# MOCHI enables discovery of heterogeneous interactome modules in 3D nucleome

**DOI:** 10.1101/542092

**Authors:** Dechao Tian, Ruochi Zhang, Yang Zhang, Xiaopeng Zhu, Jian Ma

**Affiliations:** Computational Biology Department, School of Computer Science, Carnegie Mellon University, Pittsburgh, PA 15213, USA

## Abstract

The composition of the cell nucleus is highly heterogeneous, with different constituents forming complex interactomes. However, the global patterns of these interwoven heterogeneous interactomes remain poorly understood. Here we focus on two different interactomes, chromatin interaction network and gene regulatory network, as a proof-of-principle, to identify heterogeneous interactome modules (HIMs) in the nucleus. Each HIM represents a cluster of gene loci that are in spatial contact more frequently than expected and that are regulated by the same group of transcription factor proteins. We develop a new algorithm MOCHI to facilitate the discovery of HIMs based on network motif clustering in heterogeneous interactomes. By applying MOCHI to five different cell types, we found that HIMs have strong spatial preference within the nucleus and exhibit distinct functional properties. Through integrative analysis, this work demonstrates the utility of MOCHI to identify HIMs, which may provide new perspectives on 3D genome organization and function.

## Introduction

The cell nucleus is an organelle that contains heterogeneous components such as chromosomes, proteins, RNAs, and subnuclear compartments. These different constituents form complex organizations that are spatially and temporally dynamic (Lanctôt et al., 2007; Bonev and Cavalli, 2016). Interphase chromosomes are folded and organized in three-dimensional (3D) space by compartmentalizing the cell nucleus (Cremer and Cremer, 2001; van Steensel and Belmont, 2017), and different chromosomal loci also interact with each other (Bonev and Cavalli, 2016). The development in whole-genome mapping approaches such as Hi-C (Lieberman-Aiden et al., 2009) to probing chromatin interactome has enabled comprehensive identification of genome-wide chromatin interactions, revealing important nuclear genome features such as loops (Rao et al., 2014; Tang et al., 2015), topologically associating domains (TADs) (Dixon et al., 2012; Nora et al., 2012), and A/B compartments (Lieberman-Aiden et al., 2009). Nuclear genome organization has intricate connections with gene regulation (Cremer and Cremer, 2001; Misteli, 2007). In particular, correlations between higher-order genome organization (including chromatin interactions and chromosome compartmentalization) and transcriptional activity have been demonstrated (Guelen et al., 2008; Rao et al., 2014; Chen et al., 2018).

Systems level transcriptional machinery can often be represented by gene regulatory networks (GRNs), which are dynamic among various cellular conditions (Gerstein et al., 2012; Marbach et al., 2016). GRN models the phenomena of selective binding of transcription factor (TF) proteins to *cis*-regulatory elements in the genome to regulate target genes (Davidson, 2006; Lambert et al., 2018). Transcription of Co-regulated genes in GRN can be facilitated by long-range chromosomal interactions (Fanucchi et al., 2013) and chromatin interactome has been shown to exhibit strong correlations with GRN (Kosak et al., 2007; Neems et al., 2016). Indeed, network-based representation of both chromatin interactome and GRN has been suggested to consider different subnuclear components holistically (Rajapakse et al., 2010; Chen et al., 2015). The paradigm of viewing the nucleus as a collection of interacting networks among various constituents can potentially be extended to account for other types of related interactomes in the nucleus. However, whether these interactomes, in particular chromatin interactome and GRN, are organized to form functionally relevant, global patterns remains unknown.

In this work, as a proof-of-principle, we specifically consider two different types of interactomes in the nucleus: (1) chromatin interactome – a network of chromosomal interactions between different genomic loci – and (2) a GRN where TF proteins bind to the genomic loci to regulate target genes’ transcription. Many studies in the past have analyzed the structure and dynamics of chromatin interactomes and GRNs as well as the coordinated binding of transcription factors on folded chromatin (Rao et al., 2014; Tang et al., 2015; Marbach et al., 2016; Ma et al., 2018; Cortini and Filion, 2018). However, the global network level patterns between chromatin interactome and GRN are still unclear, and algorithms that can simultaneously analyze these heterogeneous networks in the nucleus to discover important network structures have not been developed.

Here we aim to identify mesoscale network structures where nodes of TFs (from GRN) and gene loci (from both chromatin interactome and GRN) cooperatively form distinct types of modules (i.e., clusters). We develop a new algorithm, MOCHI (MOtif Clustering in Heterogeneous Interactomes), that can effectively uncover such network modules, which we call heterogeneous interactome modules (HIMs), based on network motif clustering using a 4-node motif specifically designed to reveal HIMs. Each identified HIM represents a collection of gene loci and TFs for which (1) the gene loci have higher than expected chromatin interactions between themselves, and (2) the gene loci are regulated by the same group of TFs. To demonstrate the utility of MOCHI to identify HIMs based on complex heterogeneous interactomes in the nucleus, we apply MOCHI to five different human cell types, identifying patterns of HIMs and their functional properties through integrative analysis. HIMs have the potential to provide new insights into the nucleome structure and function, in particular, the interwoven interactome patterns from different components of the nucleus. The source code of our MOCHI method can be accessed at: https://github.com/macompbio/MOCHI.

## Results

### Overview of the MOCHI algorithm

The overview of our method is illustrated in Fig. 1, with detailed algorithms described in the Methods section. Our goal is to reveal network clusters in a heterogeneous network such that certain higher-order network structures (e.g., the network motif *M* in Fig. 1A) are frequently contained within the same cluster. The input heterogeneous network in this work considers two types of interactomes: a GRN (directed) between TF proteins and target genes; and chromatin interaction network (undirected) between gene loci on the genome. For chromatin interactome, for each pair of gene loci within 10Mb, we use the “observed over expected” (O/E) quantity in the Hi-C data (we use O/E>1 as the cutoff in this work, but we found that our main results are largely consistent with different cutoffs; see Supplemental Information B.1) to define the edges in the chromatin interaction network. For GRN, we use the transcriptional regulatory networks from (Marbach et al., 2016), which were constructed by combining enrichment of TF binding sites in enhancer and promoter regions and co-expression between TFs and genes. If a TF protein regulates a gene, we add a directed edge from the TF to the gene. We then merge the chromatin interaction network and the GRN from the same cell type to form a network *G* with nodes that are either TF proteins or gene loci together with the directed and undirected edges defined above (Fig. 1B).

**Figure 1:**
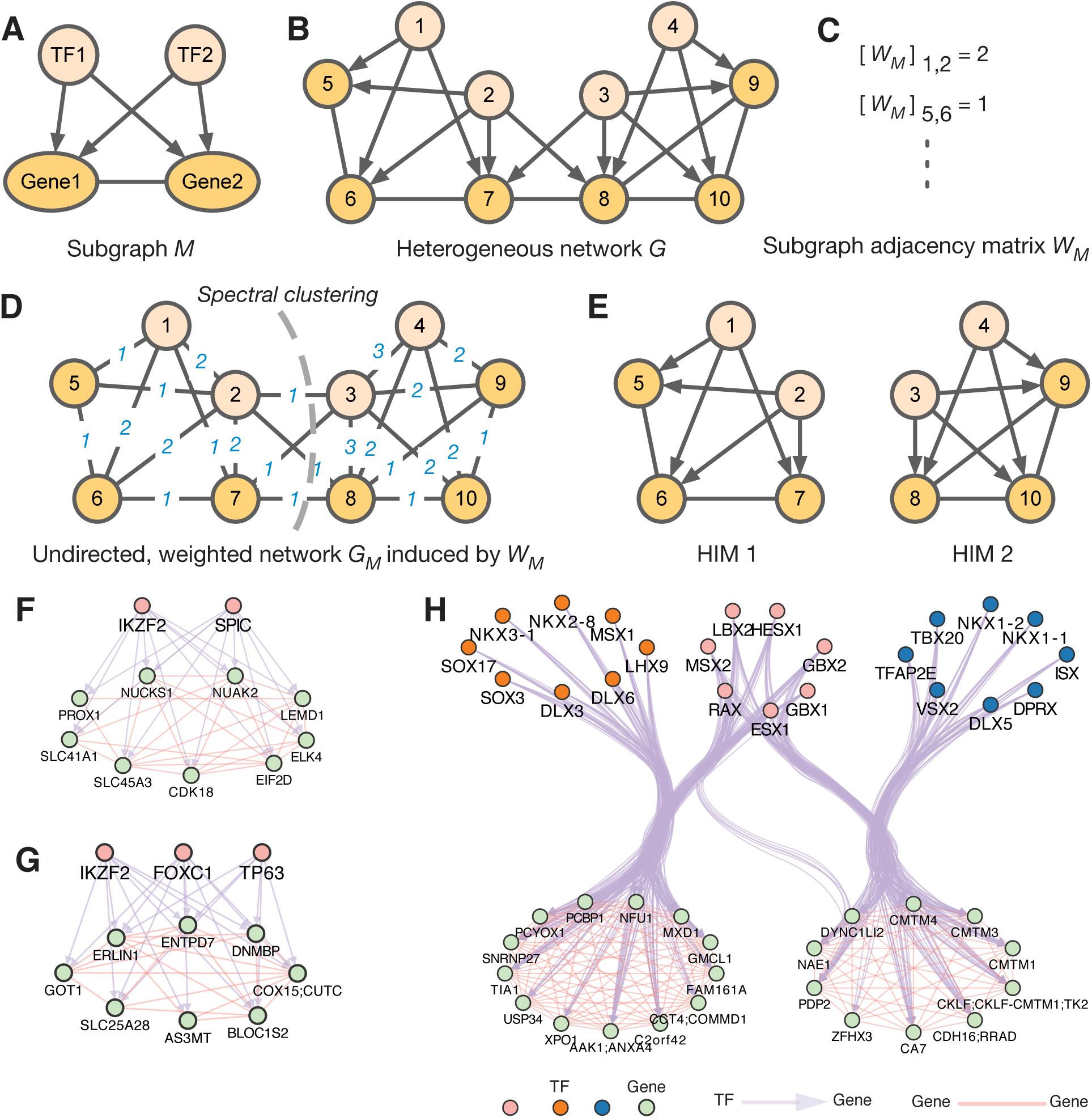
Workflow of our MOCHI algorithm and output examples of HIMs. The network has both genegene spatial proximity and TFgene regulation relationships. **(A)** A 4node motif *M* represents the smallest HIM. Here directed interaction represents a TF-gene regulation relationship, an undirected interaction represents that the two genes are spatially more proximal to each other than expected. **(B)** Given a heterogeneous network *G*, we find HIMs by minimizing the motif conductance (see Eq. 2). **(C)** We compute the subgraph adjacency matrix *W*_*M*_ with [*W*_*M*_]*_ij_* being the number of occurrences of *M* that have both nodes *i* and *j*. **(D)** The weighted network *G*_*M*_ is defined from adjacency matrix *W*_*M*_. **(E)** Spectral clustering will find clusters in *G*_*M*_. We recursively apply the method to find multiple HIMs and overlapping HIMs. **(FG)** Two HIMs as examples in GM12878. **(H)** Example of two overlapping HIMs in GM12878 sharing 7 TFs (the group with pink nodes). TFs in orange and pink nodes form one HIM with their target genes (bottom left). TFs in pink and blue nodes form another HIM with their target genes (bottom right). Note that the directed interactions from TFs to their target genes are bundled.

We specifically consider the network motif *M* which has four nodes, i.e., two gene loci and two TFs in the heterogeneous network with two genes whose genomic loci are spatially more proximal to each other (than expected) within the nucleus and that are also co-regulated by the two TFs (Fig. 1A) (see the Methods section and Supplemental Information for the justification of this motif). Our goal is to reveal higher-order network clusters based on this particular network motif. In other words, we want to partition the nodes in the network such that this 4-node network motif occurs mostly within the same cluster. Based on the motif, our MOCHI algorithm, which extends the original algorithm in (Benson et al., 2016), constructs an undirected, weighted network *G*_*M*_ (Fig. 1D) based on subgraph adjacency matrix *W*_*M*_ (Fig. 1C). We then apply recursive bipartitioning in *G*_*M*_ to find multiple clusters (Fig. 1E). We call such clusters HIMs, which, in this work, represent network structures containing the same group of TFs that regulate many target genes whose spatial contact frequencies are higher than expected. Since TFs can regulate multiple sets of genes that may belong to different clusters, different HIMs may overlap by sharing TFs. The algorithm details of MOCHI are in the Methods section.

### MOCHI identifies HIMs in multiple cell types

We applied MOCHI to five different human cell types: GM12878, HeLa, HUVEC, K562, and NHEK. The input heterogeneous network of each cell type has 591 TFs, ∼12,000 expressed genes, and ~1 million regulatory interactions (Table S1). A few examples of HIMs identified in GM12878 are shown in Fig. 1F-H, including overlapping HIMs in Fig. 1H. We found that the identified HIMs in five cell types share several basic characteristics. The number of identified HIMs ranges from 650 to 806 in different cell types, with at least 71.9% of the HIMs sharing TFs with other HIMs in each cell type. Notably, HIMs cover a majority (62.1-77.2%) of the genes in the heterogeneous networks (Table S1). For example, in GM12878, there are 591 TFs co-regulating 7,617 (69.1%) genes in 650 HIMs. The HIMs have, on average, 9-17 TFs regulating 9 genes in different cell types (Table S2). In addition, we found that the identified HIMs in different cell types share similar connections to 3D genome features (Supplemental Information B.3, Table S2).

To further assess that the genes in a HIM are indeed co-regulated by the same TF, we used the available ChIP-seq data of 26 TFs in GM12878 and K562 cells from the ENCODE project (Consortium et al., 2012). We found that for all the HIMs in GM12878 or K562 with these 26 TFs, more than half (55.85%) of them have ≥50% of their genes with corresponding TF ChIP-seq peaks within 10kb of the transcription start site, further suggesting that the genes in HIMs identified by MOCHI share regulatory TFs. In addition, MOCHI can reliably identify HIMs with different parameters in various cell types (Supplemental Information B.1). Importantly, we justified the choice of the 4-node motif *M* by showing its advantages over a triangle motif and a bifan motif (Supplemental Information B.2). The triangle and bifan motifs do not explicitly encode the co-regulation between TFs and the spatial proximity between genes. These results demonstrate that MOCHI can reliably identify HIMs across multiple cell types.

### HIMs show strong preference in spatial location relative to subnuclear structures

Next, we specifically analyzed the spatial localization of HIM in the nucleus. Recently published SON TSA-seq and Lamin B TSA-seq datasets quantify cytological distance of chromosome regions to nuclear speckles and nuclear lamina, respectively (Chen et al., 2018). In K562, which is currently the only cell type with TSA-seq data, 60.7% of the HIMs have mean SON TSA-seq score higher than 0.284 (80-th percentile of the SON TSA-seq score), suggesting that the genes in these HIMs, on average, are within 0.518*µ*m (estimated in Chen et al. (2018)) of nuclear speckles (Fig. 2A). Compared to the genes in the K562 heterogeneous network but not assigned to HIMs, the genes in HIMs have higher SON TSA-seq score and lower Lamin B TSA-seq score (*p<*2.22e-16; Fig. S1).

**Figure 2:**
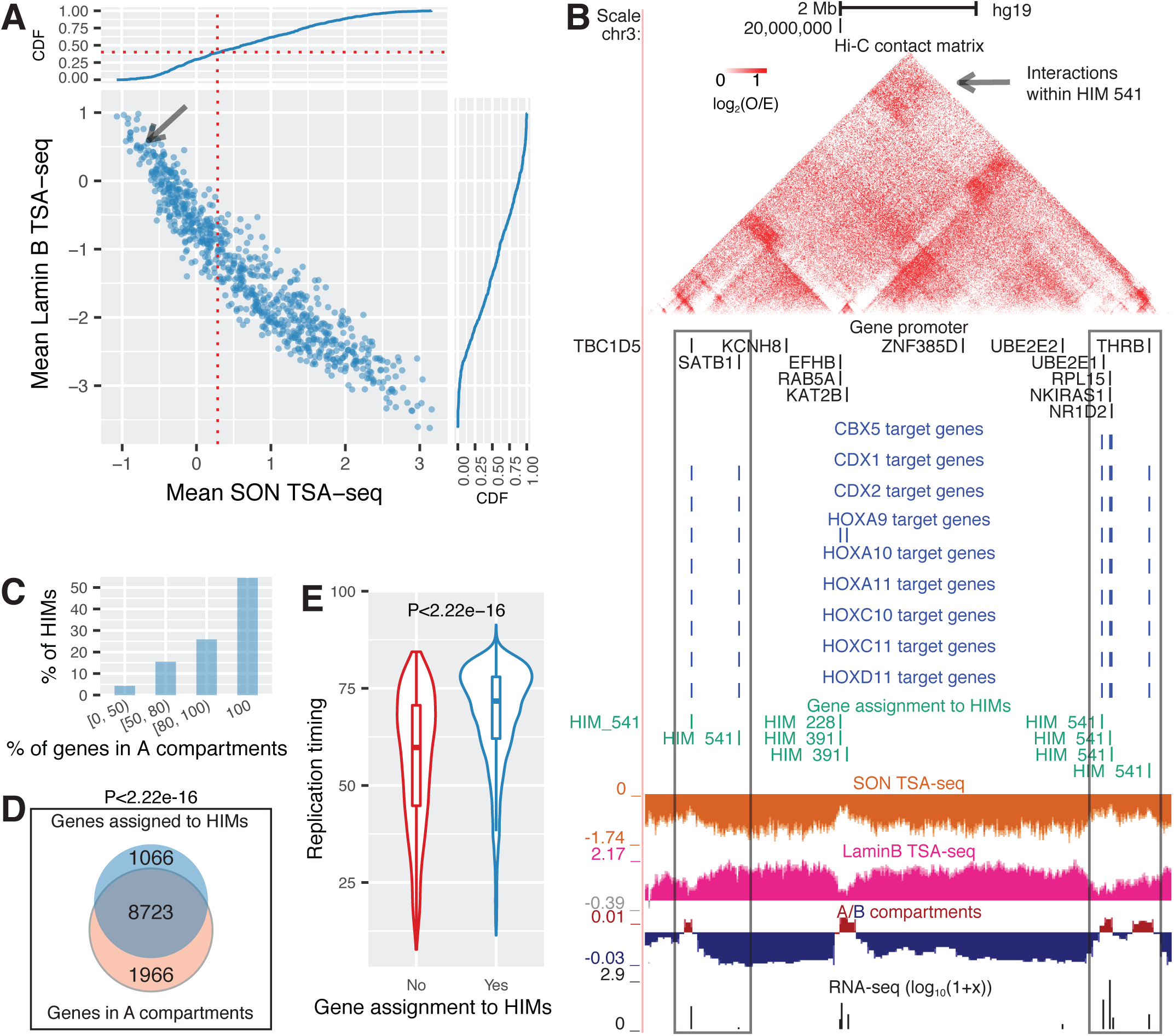
HIMs tend to be close to nuclear interior, in particular, speckles. **(A)** Scatter plot shows the mean SON TSA-seq score and mean Lamin B TSA-seq score of the genes in each HIM. Each dot represents a HIM. The curves on top and right are cumulative density functions (CDF). The red vertical dotted line represents the mean SON TSA-seq at 0.284 (approx. within 0.518 µm of nuclear speckles (Chen et al., 2018)). The black arrow points to HIM #541. **(B)** HIM #541 with low mean SON TSA-seq (pointed by the arrow in **(A)**). The heatmap shows the upper-triangle part of the Hi-C contact matrix (O/E) of the 10kb-sized bins in the chromosome region that covers the genes in the HIM. Target genes of different TFs, gene members of HIM, SON TSA-seq, LaminB TSA-seq, A/B compartments, and RNAseq signals are shown in different tracks. **(C)** Barplot shows the proportion of HIMs with a varied proportion of genes in A compartment. **(D)** Venn diagram shows that the genes assigned to HIMs are enriched in A compartment. **(E)** Violin and boxplot compare the replication timing of the genes assigned to HIMs and the other genes in the heterogeneous network of K562. Here the HIMs are identified in K562 cell line. The spatial location features of HIMs in other cell types are in Fig. S2.

We specifically looked at the HIMs that are away from the nuclear interior. Fig. 2B shows one HIM (#541) that is close to nuclear lamina (mean Lamin B TSA-seq score 0.593, mean SON TSA-seq score 0.642). This HIM has 9 TFs co-regulating 6 genes that span 6.78 Mb on chromosome 3. The Hi-C edge density (see Supplemental Information A.4) among these genes is 0.667, suggesting that these 6 genes as a group are spatially closer to each other than expected (i.e., connected with chromatin interaction). The SON TSA-seq scores of the 6 genes are low but tend to be the local maxima (i.e., small peaks within valleys), while the Lamin B TSA-seq scores are high but tend to be the local minima (i.e., small valleys within peaks), suggesting that these gene loci are localized more towards the nuclear interior than their surrounding chromatin. Five out of the 6 genes are expressed with FPKM*>*=3.4. The gene RPL15 in this HIM is a K562 essential gene (Wang et al., 2015). The TF proteins CDX1, HOXA9, and HOXA10 are involved in leukemia and hematopoietic lineage commitment provided by Genecards (Safran et al., 2010). This suggests that even though HIM #541 is a HIM away from nuclear speckle, it may play relevant functional roles in K562.

Recently, Quinodoz et al. (2018) reported that inter-chromosomal interactions are clustered around two distinct nuclear bodies as hubs, including nuclear speckles and nucleoli. By comparing with the genomic regions organized around nucleolus based on data from the SPRITE method in GM12878 (Quinodoz et al., 2018), we found that vast majority (85.4%) of the GM12878 HIMs do not have genes close to the nucleolus. Earlier work estimated that only 4% of the human genome is within nucleolus-associated domains (Németh et al., 2010). It is therefore expected that only a small number of HIMs would be close to the nucleolus. Indeed, we found that there are only 30 (4.62%) GM12878 HIMs with all their genes near the nucleoli. Notably, 16 out of these 30 HIMs have at least one TF protein located close to nucleoli according to protein subcellular locations from the human protein atlas (Thul et al., 2017). For example, HIM #267 has 4 TF regulators: ETS1, ETV6, PPARG, and PTEN, where ETV6 is known to localize to the nucleoli.

Earlier work from Hi-C data showed that at megabase resolution the interphase chromosomes are segregated into A and B compartments that are largely active and inactive in transcription, respectively (Lieberman-Aiden et al., 2009). Chromosome regions in B and A compartments have nearly identical agreements with lamina associated domains (LADs) and inter-LADs (i.e., more towards interior) (van Steensel and Belmont, 2017). Compartment A regions also replicate earlier than compartment B regions (Pope et al., 2014). We found that the genes in HIMs are preferentially in A compartments and replicated earlier across cell types. Specifically, 57.4% of HIMs have genes that are all in A compartments in K562. Only a small proportion (4.49%) of HIMs have over 50% of genes in B compartments (Fig. 2C). We found that the genes in HIMs as a whole are more enriched in A compartments, with 89.1% of them in A compartments (*p<*2.22e-16, hypergeometric test; Fig. 2D). Compartment A can be further subdivided into A1 and A2 subcompartments in GM12878 (Rao et al., 2014) at a finer scale. Among the 369 GM12878 HIMs with genes all in A compartments, 198 (53.66%) HIMs have *≥*80% of their genes in A1 subcompartments, 60 (16.26%) HIMs are in A2 subcompartments, and the rest 111 HIMs span both A1 and A2 compartments. Additionally, we found that the genes assigned to HIMs have much earlier replication timing than the other genes (*p<*2.22e-16; Fig. 2E). We also observed that the genes on the same chromosome that are in HIMs tend to have more similar replication timing as compared to the genes (on the same chromosome) that are not in HIMs (Fig. S2). These patterns can also be observed in other cell types (Fig. S2).

Taken together, these results suggest that HIMs have strong preference to localize towards the nuclear interior in active compartments with the majority of them being in proximity of the nuclear speckles and replicating earlier.

### HIMs are enriched with essential genes, super-enhancers, and PPIs

Next, we explored the functional properties of HIMs. We again grouped the genes assigned to HIMs into one set and the genes in the heterogeneous network but are not assigned to HIMs into another set. For a fair comparison, we also stratify the gene sets by chromosome number. We call these clusters merged-HIM clusters and non-HIM clusters accordingly. We first compared with the information of gene essentiality (Wang et al., 2015) (see Supplemental Information A.5). We found that genes assigned to HIMs are enriched with essential genes across all five cell types. For example, 12.7% of the genes assigned to HIMs in K562 are K562 essential genes, which is significantly higher than the proportion (7.79%) of the genes not assigned to HIMs (*p*=1.13e-12; Fig. 3A). This observation is also present across chromosomes (Fig. S3A). Across the cell types, genes assigned to HIMs consistently have higher proportions of essential genes than genes not assigned to HIMs (*p ≤*2.17e-6; Fig. S3B). Regarding gene expression level, we found that genes assigned to HIMs are more highly expressed and expressed at similar levels (Fig. 3B, Fig. S4).

**Figure 3:**
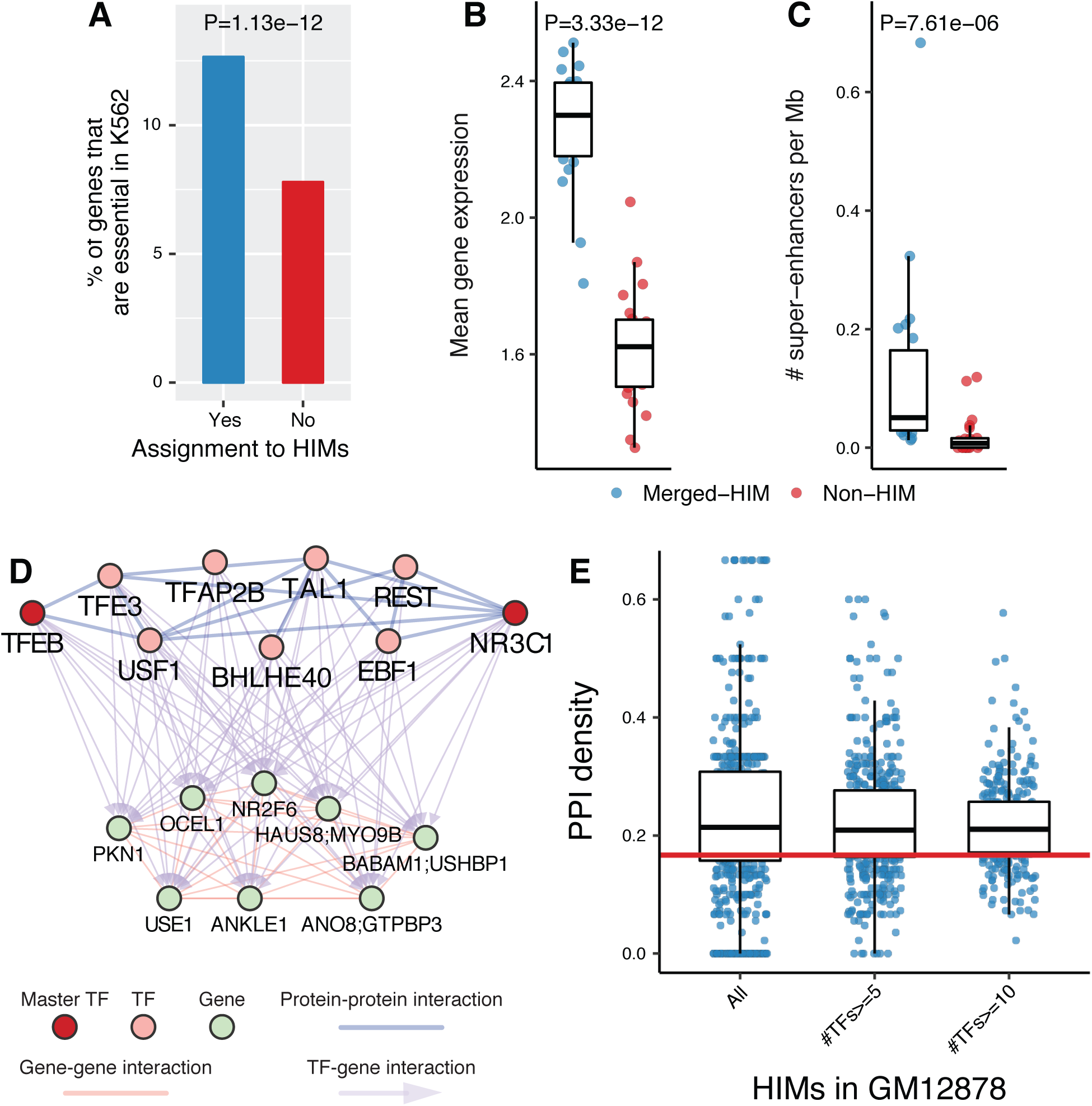
HIMs are enriched with essential genes, super-enhancers, and protein-protein interactions. **(A)** Barplots show the proportions of genes that are K562 essential genes among the genes assigned to HIMs and those not assigned to HIMs. **(B-C)** Functional properties of the genes in the identified HIMs in K562. To make a fair comparison, we stratified the genes assigned to HIMs by chromosome number and called resulted clusters as merged-HIM clusters. Similarly, we derived non-HIM clusters from the genes in the heterogeneous networks but not assigned to HIMs. Pvalues are computed by the paired twosample Wilcoxon ranksum test. **(B)** Boxplot shows the average gene expression level of the genes in a cluster. **(C)** Boxplot shows the normalized number of super-enhancers related to a cluster. **(D-E)** TFs in HIMs are enriched with protein-protein interactions (PPIs) among themselves. **(D)** One example of HIM from GM12878 cell line shows that 9 TFs in the HIM are connected by 14 PPI interactions. The sub-PPI network has a density at 0.389. The TFs NR3C1 and TFEB are master TFs in GM12878. **(E)** Boxplots show the distribution of the sub-PPI network density of the HIMs and the subsets of HIMs with at least *n* TFs, *n* = 5, 10. The medians are significantly (*p<*2.22e16) higher than the expected density (0.158, red line) of the sub-PPI networks induced by randomly sampled TFs.

Super-Enhancers are known to be associated with many cell type-specific functions (Hnisz et al., 2013). To study the connections between HIMs and super-enhancers, we computed the cluster-size normalized number of super-enhancers annotated in (Hnisz et al., 2013) that (1) have Hi-C contacts with, and (2) are close to (window size=50kb) at least one gene in each cluster. We found that HIMs are enriched with spatial contacts with super-enhancers. Specifically, the merge-dHIMs have at least 6-fold higher normalized number of super-enhancers than the non-HIMs across cell types (Fig. 3C, Fig. S5). The significant pattern is consistent with a varied window size from 20kb to 1Mb (Fig. S5).

Protein-Protein interactions (PPIs) can further stabilize TF-DNA binding of the interacting TFs (Lambert et al., 2018). We ask whether TFs in the same HIM tend to have more PPIs with each other. We computed the density of the sub-PPI network induced by the TFs in a HIM, where the PPI network is based on 591 TF proteins used in this study (see Supplemental Information A.5). We found that TFs within HIMs are enriched with PPIs among themselves as compared to random cases selected from the 591 TFs. For example, in GM12878, TFs NR3C1 and TFEB, which are master regulators (Hnisz et al., 2013), co-regulate 8 genes with the other 7 TFs proteins in a HIM (Fig. 3D). This particular sub-PPI network of the 9 TFs has 14 interactions. The density of this sub-PPI network is 0.389 which is 2.46 times higher than the average density (0.158) of the random cases. Overall, the median density of the sub-PPI networks induced by TFs in the identified HIMs in GM12878 is 0.214, much higher than the random cases (*p<*2.22e-16; Fig. 3E). This observation is also consistent in other cell types in this study (Fig. S6). We also found that the significance is not affected by a varied number of TFs across HIMs (Fig. S6).

These results suggest that the genes and TFs involved in HIMs likely perform critical roles, which are manifested by the level of gene essentiality of target genes, engagement of super-enhancers, and enrichment of PPI among TFs.

### Genes in HIMs exhibit stability and variability across cell types

To study how HIMs change across different cell types, we first focused on the assignment of genes to HIMs in different cell types. Through pairwise comparison, we found that the genes assigned to HIMs have the highest degree of overlap between GM12878 and K562 as compared to the other cell types, which is consistent with the fact that both GM12878 and K562 are from human hematopoietic cells (Fig. S7A). Comparisons among all five cell types showed that 3,025 genes are consistently assigned to HIMs, accounting for 30.91% to 40.06% of genes that are in the HIMs in each cell type (Fig. 4A). In contrast, only a small fraction (*≤* 5.93%) of genes are uniquely assigned to the HIMs in each cell type. For example, out of the 8,034 genes in the GM12878 HIMs, only 344 (4.28%) genes are not assigned to HIMs in other cell types (Fig. 4A).

**Figure 4:**
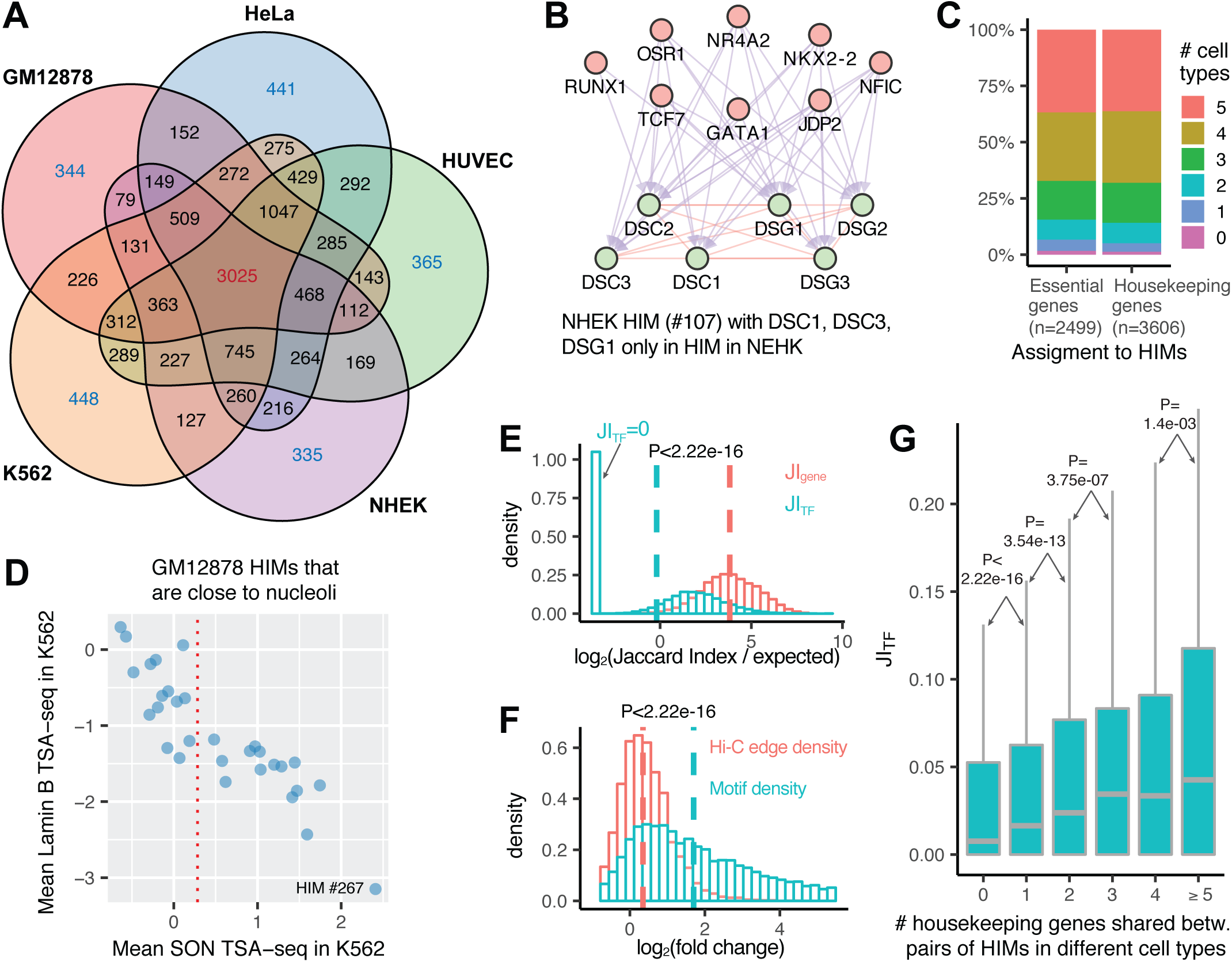
HIM comparisons in terms of genes and TFs across the cell types. **(A)** Venn diagram shows the assignment of genes in HIMs across five cell type. Numbers in each facet represent the gene number in each possible logic intersection relationship across five cell types. **(B)** A NHEK HIM with 3 genes only assigned to HIMs in NHEK. All of its genes are involved in keratinization pathway. Here the top and bottom nodes are the TFs and genes in the HIM, respectively. **(C)** Barplot shows the assignment of essential genes and housekeeping genes to HIMs across five cell types. **(D)** Scatter plot shows the mean SON TSA-seq and Lamin B TSA-seq scores (in K562 (Chen et al., 2018)) of the 30 GM12878 HIMs that are inferred as close to nucleoli in GM12878 (Quinodoz et al., 2018). The red vertical dotted line represents the mean SON TSA-seq score at 0.284. **(E)** The logtransformed ratio of Jaccard index on the genes/TFs between paired HIMs from different cell types over the expected Jaccard index between random control sets. **(F)** Fold changes of motif *M* density and Hi-C edge density of each HIM between the cell type it is identified and another cell type. Here a vertical dash line represents the median of a variable. **(G)** Boxplots show the distribution of Jaccard index on the TFs of paired HIMs with different numbers of shared housekeeping genes.

The genes consistently and uniquely assigned to HIMs are enriched with distinct functional terms using DAVID (Huang et al., 2008) (Table S5). The genes consistently assigned to HIMs are strongly enriched with functions related to essential cellular machinery, whereas the genes uniquely assigned to HIMs in a particular cell type are enriched with more cell type-specific functions. An example is NHEK HIM #107 (Fig. 4B). Among the 6 genes in this HIM, DSC1, DSC3, DSG1 are not assigned to HIMs in the other cell types. These 6 genes are involved in the keratinization pathway based on GeneCards (Safran et al., 2010). We further assessed the assignment of housekeeping genes (Eisenberg and Levanon, 2013) and essential genes to HIMs. We found that for both sets of genes, the majority (*≥* 84%) of them are assigned to HIMs consistently or in at least 3 out of the 5 cell types (Fig. 4C), suggesting that the genes with crucial functions tend to form spatial clusters across multiple cell types.

We next analyzed the variability of HIMs in terms of spatial proximity to subnuclear compartments. We found that 15 out of the 30 HIMs close to nucleoli in GM12878 (based on the data from (Quinodoz et al., 2018)) have mean SON TSA-seq score *≥*0.284 in K562 (based on the data from (Chen et al., 2018)) (Fig. 4D), in other words, these HIMs are involved in a change of spatial position from nucleoli to speckle between GM12878 and K562. One notable example is HIM #267 in GM12878 which has the highest mean SON TSA-seq score (2.41) in K562. Interestingly, the 10 genes (in HIM #267 in GM12878) together with another 8 genes form a new HIM (#628) in K562. This GM12878 HIM #267 has four TFs: ETS1, ETV6, PPARG, and PTEN. On the other hand, the K562 HIM #628 has four different TFs: KLF4, NFKB1, STAT3, and WT1, where KLF4, STAT3, and WT1 are known to be involved in the progression of leukemia.

To compare the detailed membership changes of HIMs across cell types, we computed Jaccard indices, denoted by JI*_TF_* and JI*_gene_*, of the TF members and gene members between HIMs from two different cell types, respectively. We found that the gene members undergo a moderate change from one cell type to another, whereas the TF members change at a much higher rate. JI*_gene_* has a median of 0.096 and it is higher than expected JI*_gene_* between random gene sets while controlling the set size and chromosome number (median ratio=14.12, Fig. 4E). On the other hand, JI*_T F_* has a median of 0.017 and it is close to expected JI*_T F_* between randomly selected control TF sets (median ratio=0.878, Fig. 4E). There are at least two phenomena jointly contributing to these observations. First, the Hi-C interaction networks and GRNs are highly cell type-specific, as 66% Hi-C interactions and 31.4% GRN interactions only exist in one cell type (Table S3). Second, given a HIM identified in a cell type, the motif *M* density of the HIM (see Supplemental Information A.4) has higher fold change than the Hi-C edge density of the HIM in another cell type (*p<*2.22e-16; Fig. 4F). In other words, the co-regulation relationships of the TFs on the genes in HIMs change more often across cell types than the spatial proximity relationships between the gene loci. However, we observed that if HIMs from two different cell types share a higher number of housekeeping genes, they tend to have a higher JI*_T F_* (Fig. 4G). We found a similar pattern for essential genes (Fig S7B).

### Conserved and cell type-specific HIMs have distinct properties

Motivated by the gene membership dynamics of HIMs across cell types, we further classified HIMs into conserved and cell type-specific HIMs. For HIMs in a given cell type, we call a HIM conserved if it shares a significantly high proportion of genes (JI*_gene_ ≥* 1/3, *p≤*0.001, Bonferroni adjusted hypergeometric test) with at least one HIM in other cell types (i.e., the HIM is recurrent). Note that JI*_gene_ ≥*1/3 represents that two equal-sized gene sets share than half of their genes. The rest are called cell type-specific HIMs. As a result, 40.69-47.38% of the identified HIMs in each cell type are cell type-specific HIMs. Fig. 5 shows a cell type-specific HIM, HIM #712, in K562 and its changes in other cell types. This HIM covers 9 genes on chromosome 11. These genes spatially contact each other at higher frequencies than expected (Fig. 5A) and are co-regulated by TF protein BCL6B and CPEB1 in K562 (Fig. 5B). In other cell types, at most 4 out of the 9 genes are assigned to HIMs (Fig. 5C). We found that this HIM has K562-specific chromosomal structures and functional annotations. The genomic region covering the genes in the HIM is in A compartment in K562 but switches to B compartment in other cell types (Fig. 5D). One nearby upstream region is annotated as a superenhancer only in K562 (Hnisz et al., 2013) (Fig. 5E). Many sites are annotated as transcriptionally active states, such as enhancers, promoters, or transcribed states in K562, but not in other cell types based the results from ChromHMM (Ernst and Kellis, 2012) (Fig. 5F). The genes MRPL16, OSBP, and PATL1 are essential genes in K562. This example demonstrates that the K562-specific HIM has specific chromatin organization and biological functions.

**Figure 5:**
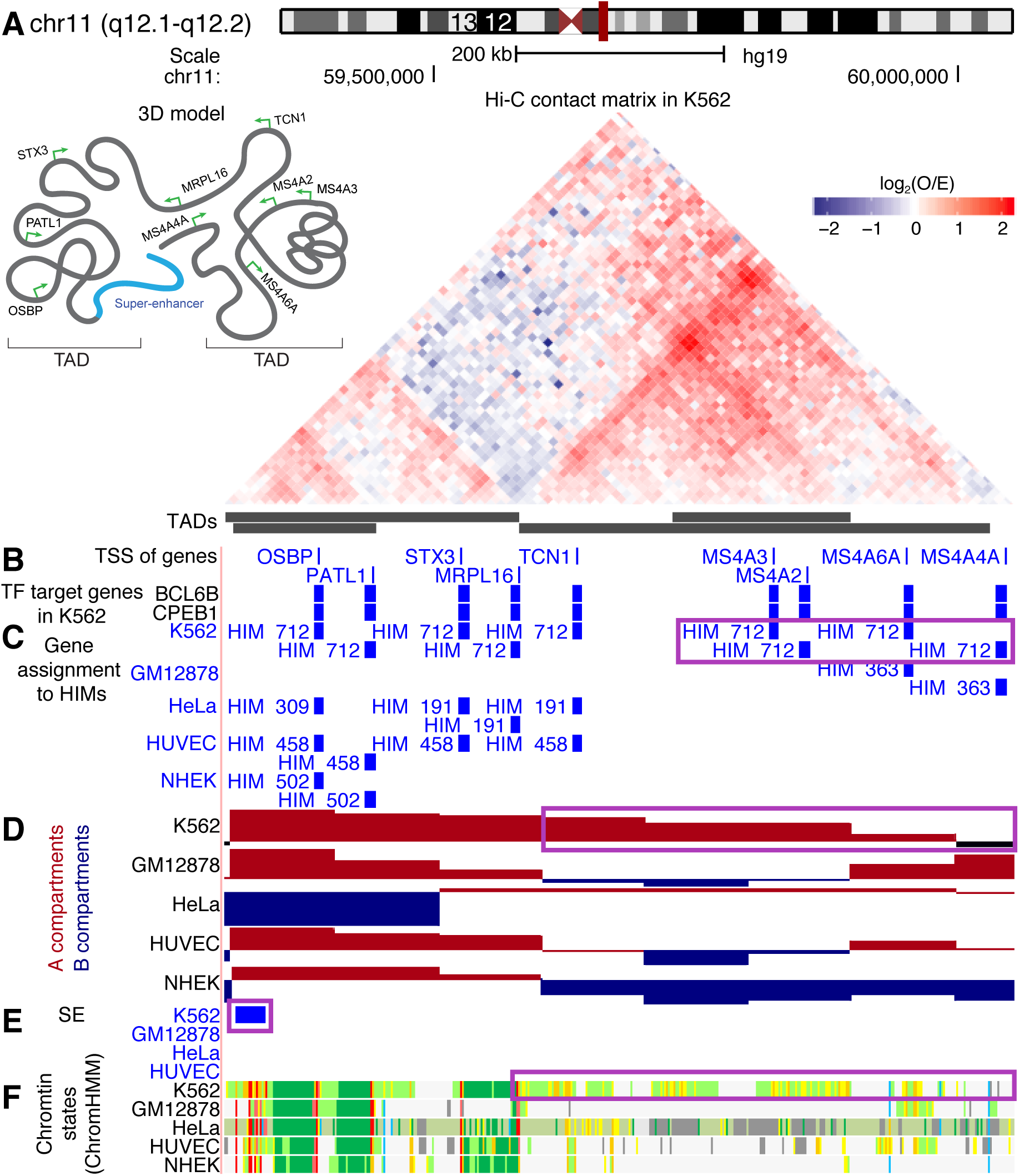
A K562 specific HIM with K562 specific chromatin interactome and functional annotations. **(A)** The 45 degree rotated upper triangle part of the contact matrix between the 10kb-sized bins in a chromosome region in K562. The region is segregated into 4 nested TADs. **(B)** Thin bars represent the transcriptional start sites (TSSs) of the genes that are in the heterogeneous networks. THi-Ck bars represent the genes that are regulated by BCL6B or CPEB1 in K562. **(C)** The assignment of the genes to HIMs in K562 and the other cell types. **(D)** The assignment of the bins to A/B compartments. **(E)** The regions that are annotated as super-enhancers (SE). **(F)** The chromatin states inferred by ChromHMM based on multiple histone modification marks, where red and purple colors represent promoters, orange and yellow stand for enhancers, green represents transcribed regions, gray represents other types of regions such as repressed regions.

Overall, we found that the conserved and cell type-specific HIMs have distinct properties of interactomes across cell types. Compared to cell type-specific HIMs, conserved HIMs exhibit stronger clustering features with higher Hi-C edge density, higher GRN edge density, and higher motif *M* density (*p ≤*8.15e-4; Fig. S8). Also, conserved HIMs tend to be closer to the nuclear interior with a higher proportion of their genes in A compartment, and their genes replicate earlier and synchronously (Fig. S8). Moreover, we found that conserved HIMs and cell type-specific HIMs tend to have large differences in gene expression level and cell type-specific genes. The conserved HIMs have higher mean gene expression level than the cell type-specific HIMs in 3 cell types except for NHEK and HUVEC (*p<*0.05; Fig. S9). On the other hand, cell type-specific HIMs have a higher proportion of cell type-specific genes (*p≤*0.02; see Supplemental Information A.5) than conserved HIMs across five cell types (Fig. S9). Taken together, our results demonstrate that conserved and cell type-specific HIMs, in general, have distinct network properties, spatial location preference, and functional characteristics.

## Discussion

To better understand the heterogeneous nature of different components in the nucleus, new computational models are needed to consider different types of molecular interacting networks. In this work, we developed MOCHI to specifically consider two types of different interactomes in the cell nucleus: (1) a network of chromosomal interactions between different gene loci, and (2) a GRN where TF proteins bind to the genomic loci to regulate target genes. MOCHI is able to identify network structures where nodes of TFs (from GRN) and gene loci (from both chromatin interactome and GRN) cooperatively form distinct network clusters, which we call HIMs, by utilizing a new motif clustering framework for heterogeneous networks. To the best of our knowledge, this is the first algorithm that can simultaneously analyze these heterogeneous networks within the nucleus to discover important network structures and properties. By applying MOCHI to five different human cell types, we made new observations to demonstrate the biological relevance of HIMs in 3D nucleome.

Our method has multiple methodological contributions. We further extended the motif conductance clustering method (Benson et al., 2016) to find overlapping HIMs in heterogeneous networks. Our study shows the utility of our new algorithm to identify HIMs based on complex heterogeneous molecular interactomes. In addition, our method can be further modified to identify other types of potentially interesting HIMs in heterogeneous networks by replacing the 4-node motif *M* with relevant motifs, especially when additional types of interactomes are included. For example, in addition to considering chromatin interactions and protein-DNA interactions as we did in this work, it would be of interest to incorporate other types of relevant interactomes in the nucleus, such as the RNA-chromatin interactome (Nguyen et al., 2018).

How can we explain the formation of HIMs? In Fig. 6, we illustrate a possible model of HIMs within the nucleus. HIMs (light pink domains) are toward the interior with a group of interacting TFs and chromatin loci. The set of TFs in a HIM cooperatively regulate target genes, which also have higher contact frequency than expected. Note that this is conceptually consistent with recently reported colocalization of TF pairs (Ma et al., 2018). Some of these TF clusters may be related to the localization preferences of TF proteins in nuclear compartments, such as nuclear speckles that are enriched with various transcriptional activities (Spector and Lamond, 2011; Chen et al., 2018). Indeed, we found that the majority of the identified HIMs are close to nuclear speckles. The definitions of HIMs may also have intrinsic connections with the emerging findings on the mechanism of nuclear subcompartment formation, where TFs and their potential regulating genes/chromatin are trapped by localized liquid-like chambers through the phase separation (Shin and Brangwynne, 2017; Hnisz et al., 2017). Evidence has been shown that phase separation can help explain the formation of superenhancer mediated gene regulation (Hnisz et al., 2017; Boija et al., 2018). From our analysis, we found that genes assigned to HIMs are enriched with contacts with super-enhancers. The genes consistently assigned to HIMs are enriched with essential biological processes related to chromosomal organization and transcription. However, the detailed formation mechanisms for HIMs, which may involve both *cis* elements and *trans* factors, remain to be investigated. It would also be important to delineate the different roles of both different TFs and different genes in forming the HIMs, as some of them may be necessary and others may be redundant for the stability of HIMs. In addition, more experimental data are needed to further evaluate the functional significance of HIMs. For example, although we observed connections between HIMs and 3D genome organization features, the intricate functional relevance among these different higher-order nucleome units that jointly contribute to gene regulation in different cellular conditions is yet to be revealed. Nevertheless, HIMs may become a useful type of nuclear genome unit in integrating heterogeneous nucleome mapping data, which has the potential to provide new insights into the interplay among different constituents in the nucleus and their roles in 3D nucleome structure and function.

**Figure 6:**
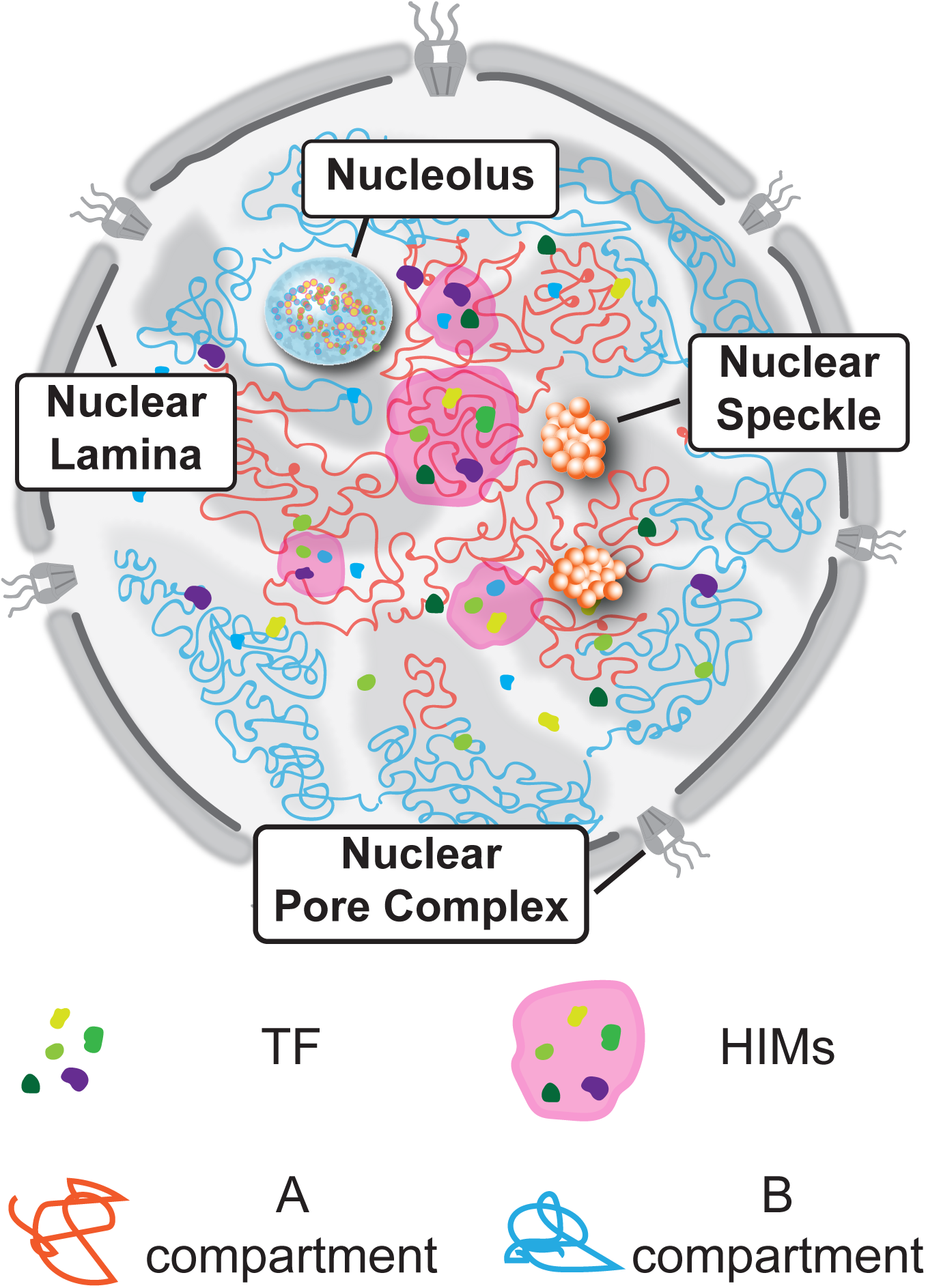
A possible model of HIMs within the nucleus.

## Methods

### Brief introduction to homogeneous network clustering by motif conductance

We first review higher-order network clustering method that can identify a cluster of nodes *S* based on motif conductance (defined below). We then introduce our algorithm MOCHI in the next subsection. Let *G* be an undirected graph with *N* nodes and *A* be the adjacency matrix of *G*. [*A*]*_ij_ ∈*{0,1} represents the connection between nodes *i* and *j*. The *conductance* of a cut 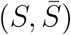, where *S* is a subset of the nodes is defined as:

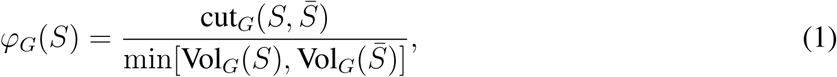

where cut 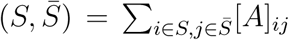 is the number of edges connecting nodes in *S* and 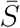. vol_*G*_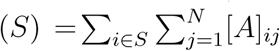 is the sum of the node degree in *S*. Moreover, the conductance of the graph *G, φ^G^*, is defined as min*_S_ φ_G_*(*S*). The *S* that minimizes the function is the optimal solution. Finding the optimal *S* is NP-hard, but spectral methods such as Fiedler partitions can obtain clusters effectively (Chung, 2007). Recently, the conductance metric is generalized to motif conductance (Benson et al., 2016; Tsourakakis et al., 2017), where a motif refers to an induced subgraph. The motif conductance computes cut*_G_* 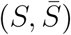) and Vol*_G_*(*S*) based on a chosen *n*-node motif. When *n* = 2, the motif is an interaction that reduces the motif conductance to conductance in Eq. (1). When *n ≥* 3, the motif conductance may reveal new highe-rorder organization patterns of the network (Benson et al., 2016). A more recent network clustering method that incorporates network higher-order structures has been developed in the setting of hypergraph clustering (Li and Milenkovic, 2017), which includes the motif conductance as a special case. However, one key limitation of the aforementioned methods is that they cannot identify overlapping clusters, which is a crucial feature of the heterogeneous networks that we want to achieve in this work.

### MOCHI – Higher-Order network clustering to identify HIMs in a heterogeneous network

We developed a higher-order network clustering method based on network motif to identify overlapping HIMs in a heterogeneous network by extending the approach in (Benson et al., 2016). We call our method MOCHI (MOtif Clustering in Heterogeneous Interactomes). We illustrate the workflow of MOCHI in Fig. 1. First, we select a specific heterogeneous 4-node network motif *M* (Fig. 1A). In *M*, two nodes are TFs and the other two nodes are genes. Both TFs regulate the two genes and the two genes are spatially more proximal to each other than expected. The motivation for choosing the subgraph *M* is that it is the building block of HIMs given that our goal is to discover a group of genes that contact with each other more frequently than expected and are regulated by the same set of TFs. As compared to simpler motifs (e.g., 3-node motif where one node is TF), our 4-node motif defined here has the advantage of simultaneously considering a pair of genomic loci that interact with each other higher than expected and that are co-regulated by the same pair of TFs.

Conceptually, our method searches for HIMs with two goals. The TFs and genes in the same HIM should be involved in many occurrences of *M.* Additionally, HIM should avoid cutting occurrences of *M*, where a cut of occurrences of *M* means that only a subset of TFs and genes in the occurrences of *M* are in the HIM node set. More formally, our method aims to find HIMs with the node set *S* that minimizes the motif conductance

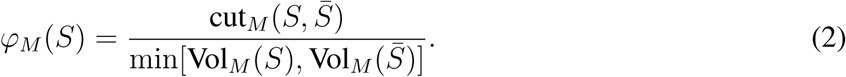

We first introduce some notations before we explain *φ*_*M*_ (*S*). Let *G* be the given heterogeneous network (e.g., Fig. 1B). Let 𝕄 be the set of occurrences of the motif *M* in *G*. For simplicity and without confusion, we also denote an occurrence of the motif *M* as *M.* Let *V*_*M*_ be the vertex set of the 2 TFs and 2 genes in *M ∈*𝕄. In Eq. (2), cut*_M_* 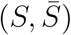 is the number of occurrences of the subgraph *M* that are cut by *S*. Formally,

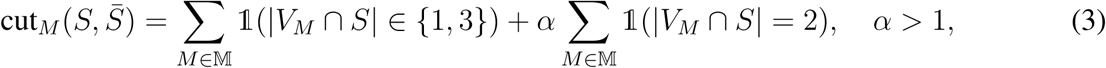

where 1 is an indicator function. Here, cut_*M*_ 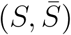 distinguishes the number of nodes of the 4-node motif *M* being assigned to *S* and 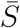. Specifically, it adds a higher penalty for the cut to the cases where two nodes in *M* are assigned to *S* and two nodes are assigned to *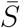*, as compared to the case where one node or three nodes are assigned to *S*, by letting *α >* 1 in Eq. (3). This is because the 1-vs-3 split could still keep interaction information from both GRN and chromatin interaction network, and the 2-vs-2 split will lose either of the information. We show that when *α* = 4*/*3 in Eq. 3 the clustering results would be near optimal (Supplemental Information A.3). Thus, *α* is set to 4*/*3 in this paper. Vol*_M_* (*S*) is the number of nodes in the occurrences of *M* that are in *S*, which is defined as:

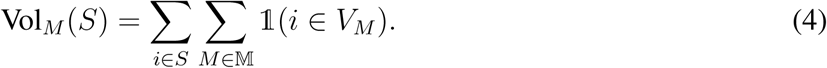

Similarly, we define the subgraph conductance of the graph *G* based on motif *M*, 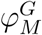 as min*_S_ φ_M_* (*S*). In the following procedures of the algorithm, we show that the motif conductance is equivalent to the normal conductance in a projection of the graph by calculating the subgraph adjacency matrix. Thus, finding the set *S* that achieves the minimum subgraph conductance is also NP-hard, following that it is NP-hard to find the minimal *φ*_*G*_(*S*). We describe our algorithm MOCHI to find HIMs that approximate the solution.

#### 1 – Calculate subgraph adjacency matrix W_M_ (*G*)

We first calculate the subgraph adjacency matrix *W*_*M*_ (*G*) by

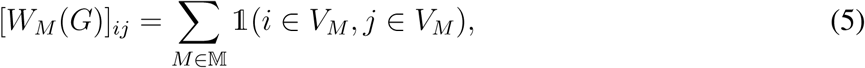

where [*W*_*M*_ (*G*)]_*ij*_ is the number of occurrences of the subgraph *M* in *G* that cover both *i* and *j* (see example in Fig. 1C). For example, if both *i* and *j* are TFs, [*W*_*M*_]*_ij_* reflects the number of paired gene loci that are spatially more proximal to each other than expected and that are also co-regulated by TFs *i* and *j*. If both *i* and *j* are genes, [*W*_*M*_]*_ij_* = 0 if *i* and *j* are not more spatially proximal to each other than expected. Otherwise, [*W*_*M*_]*_ij_* is the number of paired TFs that co-regulate *i* and *j*. Generally, *W*_*M*_ (*G*) is symmetric and [*W*_*M*_ (*G*)]*_ij_ ≥*0. Thus *W*_*M*_ (*G*) can be viewed as the adjacency matrix of an undirected weighted network. Let *G*_*M*_ denote the network with *W*_*M*_ (*G*) as the adjacency matrix (see Fig. 1D for example). It is important to note that there are genes or TFs that may not be in any occurrence of *M*, which would lead to zero vectors in the corresponding rows and columns in *W*_*M*_ (*G*). These singleton nodes in *G*_*M*_ would be removed before the next step.

#### 2 – Apply Fiedler partitions to find a cluster in G_M_

We utilize Fiedler partitions similar to the algorithm in (Benson et al., 2016) to find a cluster *S* in graph *G*_*M*_, where *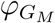* is close to the global optimal conductance of the graph: *φ*(*G*_*M*_). Recall that *φ*(*G*_*M*_) is the minimum of *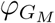* over all possible sets *S*_1_. The method is described as follows:

- Calculate the normalized Laplacian matrix of *W*_*M*_ (*G*):

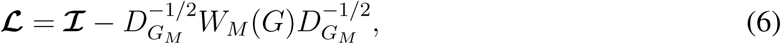

whereℐ is a identity matrix, 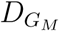 with 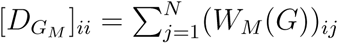 is the diagonal degree matrix of *G*_*M*_.

- Calculate the eigenvector *v* of the second smallest eigenvalue of *ℒ*.
- Find the index vector (*α*_1_,*…*, *α*_*N*_), where *α*_*k*_ is the *k*-th smallest value of 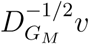.
- 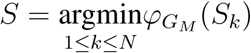 where *S*_*k*_ = {*α*_1_,*…*, *α*_*k*_}.

The sets *S* and 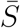 are two disjoint clusters for the heterogeneous network *G*.

#### 3 – Apply recursive bipartitioning to find multiple HIMs

We then utilize recursive bipartitioning to find multiple HIMs. We use a very different strategy than the one in (Benson et al., 2016) to select which cluster to split at each iteration, in order to specifically allow overlapping motif clusters (HIMs) with shared TFs. At each iteration, we split one HIM into 2 child HIMs. After iteration *𝓁 –* 1, there are *𝓁* HIMs: *S*_1_, *S*_2_,*…*, *S*_*𝓁*_.

At next iteration *𝓁*, one HIM *S*_*k*_ is selected if the graph it forms, *G*_*k*_, has the lowest subgraph conductance value 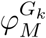 among 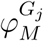, 1 <u>*≤ j*≤</u> *𝓁*. We set a threshold *t*_*1*_ for 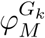. If 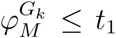, *S* will be split into two child HIMs *S*_*k*_(*c*) and 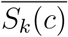 by treating the induced heterogeneous subnetwork as a new network *G*_*k*_ and repeating Steps (1) and (2) for graph *G*_*k*_. However, if the partition of graph *G*_*k*_ would lead to zero motif occurrence in either of its child graphs, we would stop partitioning this graph, add a large enough penalty value to its conductance value (to make sure it would not be selected to partition again), and move on to the next iteration. Otherwise, when 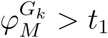, the recursive bipartitioning process will stop as all the HIM’s subgraph conductance value passes the threshold.

#### 4 – Find overlapping HIMs

Finally, we reconcile the HIMs from the clustering history tree to find overlapping HIMs. This step is added because the HIMs after Step (3) share no TFs. To reconcile the results, we first trace back the ancestral HIMs up to certain generations for each HIM based on the conductance value of its ancestor 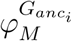, where *i* = {1, 2, 3*…*} denotes for the ‘parent’, ‘grandparent’ of the HIM. We trace along the tree until 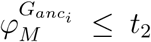, where *t*_*2*_ denotes another threshold. Clearly, *t*_*2*_ has to be smaller than *t*_*1*_ to make this process practical. Next, we pool together the TFs from the HIM and from its ancestor HIMs. We sequentially remove pooled TFs from the HIM according to their contribution to the number of occurrences of the subgraph *M* based on the graph this HIM represents, and stop this process when removing certain TF would cause a large decrease in the number of subgraphs.

#### Pseudocode and complexity of our algorithm

The pseudocode of our MOCHI algorithm is presented in the Supplemental Information A.1. The runtime of the algorithm is *O*(*t*^2^*c*^2^), where *t* and *c* (*t ≪ c*) are the number of TFs and the number of gene loci in the input heterogeneous network, respectively (detailed analysis in Supplemental Information A.2).

### Summary of the algorithm

Given a heterogeneous network from chromatin interactome network and GRN, our algorithm MOCHI will identify multiple and overlapping HIMs, which represent clusters of genes and TFs where the genes are interacting more frequently than expected and are also co-regulated by the same set of TFs. MOCHI has a few key differences as compared to the subgraph conductance method in (Benson et al., 2016). First, the input of our algorithm is a heterogeneous network with different types of nodes (TFs and gene loci), which are treated differently, while the input network for the method in (Benson et al., 2016) is rather homogeneous. Second, the algorithm in (Benson et al., 2016) will not explicitly identify multiple overlapping clusters. In MOCHI, we further developed a recursive bipartitioning method to find multiple HIMs that may overlap. Specifically, we selected a HIM to split if it has the smallest motif conductance among the HIMs at each interaction. In other words, we split the HIM that has the clearest pattern of multiple clusters. HIMs with overlapping TFs will be split in the late stage of iterations, and the overlapping information is encoded in the clustering history tree.

The recent method on hypergraph clustering (Li and Milenkovic, 2017) can be applied to identify non-overlapping HIMs where a hyperedge is defined as the motif *M.* However, similar to the method in (Benson et al., 2016), it was not designed to identify overlapping clusters, i.e., the method would not be able to find multiple overlapping HIMs that we would need. Our method also has clear differences as compared to previous works on multi-layer network clustering (see review in (Kivelä et al., 2014)). First, the inputs are different. A multi-layer network typically has only one type of nodes and different types of interactions connecting nodes within the same layer and between layers. The heterogeneous network in this work has different types of nodes (TF proteins and gene loci) and also edges. Previous multi-layer network clustering methods are not directly applicable to identify HIMs. Second, the outputs are different. The majority of multi-layer network clustering methods aim to find clusters that are either consistently observed across multiple layers or observed only in a specific layer, which are conceptually different from HIMs.

## Acknowledgement

This work was supported in part by the National Institutes of Health Common Fund 4D Nucleome Program grant U54DK107965 (J.M.), National Institutes of Health grant R01HG007352 (J.M.), and National Science Foundation grant 1717205 (J.M.). The authors would like to thank Bas van Steensel and members of Jian Ma’s laboratory (Ben Chidester, Tianming Zhou, Kyle Xiong, and Yang Yang) for helpful comments to improve the manuscript.

## Author Contributions

Conceptualization, J.M.; Methodology, D.T., R.Z., and J.M.; Software, D.T., R.Z.; Investigation, D.T., R.Z., Y.Z., X.Z., and J.M.; Writing – Original Draft, D.T., R.Z., and J.M.; Writing – Review & Editing, D.T. and J.M.; Funding Acquisition, J.M.

