## Supplemental Information for "MOCHI enables discovery of heterogeneous interactome modules in 3D nucleome"

#### A Supplementary Methods

##### A.1 Pseudocode for the MOCHI algorithm

**Algorithm 1** MOCHI

**Require:** Original graph $G_0$, motif $M$, threshold $t_1$.
**Ensure:** Network motif based clusters

**function** ITERATIVE SPECTRAL CLUSTERING($G_0, M, t_1$)
  $W_M(G_0) \leftarrow$ Motif adjacency matrix for $G_0$ based on motif $M$
  $S_0, \bar{S}_0, Score_0 =$ SPECTRAL CLUSTERING($W_M(G_0), N$)
  $L \leftarrow \{G_0\}$

  **while** $\exists G_i \in L$ such that $Score_i < t_1$, **do**
    $G_k \leftarrow \text{argmin}_i Score_i$ The graph with the lowest corresponding score
    $G_{S_k}, G_{\bar{S}_k} \leftarrow$ Graph for node set $S_k, \bar{S}_k$, respectively
    $W_M(G_{S_k}), W_M(G_{\bar{S}_k}) \leftarrow$ Motif adjacency matrix for $G_{S_k}, G_{\bar{S}_k}$
    Drop all-zero rows and columns in $W_M(G_{S_k}), W_M(G_{\bar{S}_k})$ and corresponding nodes in $G_{S_k}, G_{\bar{S}_k}$
    $N_{S_k}, N_{\bar{S}_k} \leftarrow$ Node size of $G_{S_k}, G_{\bar{S}_k}$, respectively
    $S_{S_k}, \bar{S}_{S_k}, Score_{S_k} =$ SPECTRAL CLUSTERING($W_M(G_{S_k}), N_{S_k}$)
    $S_{\bar{S}_k}, \bar{S}_{\bar{S}_k}, Score_{\bar{S}_k} =$ SPECTRAL CLUSTERING($W_M(G_{\bar{S}_k}), N_{\bar{S}_k}$)
    $L \leftarrow \{..., G_{k-1}, G_{k+1}, ..., G_{S_k}, G_{\bar{S}_k}\}$
  **end while**
**end function**

**function** SPECTRAL CLUSTERING($W_M, N$)
  $D \leftarrow$ Diagonal Matrix$_{(1:N \times 1:N)}$ given by $D_{ii} = \sum_{j=1}^{N} (W_M)_{ij}$
  $L \leftarrow D^{-\frac{1}{2}}(D - W_M)D^{-\frac{1}{2}}$
  $v = \{v_1, v_2, ..., v_N\} \leftarrow$ Eigenvector of $L$
  $v_k \in v \leftarrow$ Eigenvector of the second smallest eigenvalue
  $O \leftarrow D^{-\frac{1}{2}} v_k$
  $\alpha_i \leftarrow$ Index of the $i$-th smallest value in $O$
  $S \leftarrow \text{argmin}_k \varphi_{G_M}(S_k)$, where $S_k = \{\alpha_1, ..., \alpha_k\}$
  $Score \leftarrow \varphi_{G_M}(S)$
  **return** $S, \bar{S}, Score$
**end function**

##### A.2 Computational complexity

Here we analyze the computational complexity of MOCHI. In practice, the most time-consuming step would be the construction of the motif adjacency matrix $W_M$ and the calculation of the eigenvector for the normalized Laplacian matrix. Although in general for eigenvalue decomposition of a matrix size of $N \times N$, the runtime would be $O(N^3)$, using fast symmetric diagonally dominant solvers for Laplacian matrix, we can reach near linear time for this process (Kelner et al., 2013). Therefore, in the rest of this section, we will only discuss the computational complexity of the matrix construction part.

Intuitively, for a 4-node motif, we can calculate $W_M$ by checking every combination of 4 nodes in the graph, and has the complexity of $O(N^4)$, where $N$ is the number of nodes in the graph. However, here since we only deal with a special 4-node motif that consists of 2 different types of nodes, which can be treated as a combination of two specific 3-node motifs (one TF regulates two genes), if we use $T$ for the TF nodes in the graph, and $C$ for the chromatin loci nodes, and $t, c$ for the size of these nodes, respectively, we can derive the runtime as follows. For $W_{M_{ij}}$ in the motif adjacency matrix, where $i \in C, j \in C$, it is equivalent to finding the number $n_{3_{ij}}$ of specific 3-node motif that $i$ and $j$ share, then using the combination number to get the number of 4-node motif $n_{4_{ij}} = \binom{n_{3_{ij}}}{2}$. For the search of triangles, given $i$ and its neighbor $j$, we sum up all the TF nodes that they share, which gives us the complexity of $O(tc^2)$. For $W_{M_{ij}}$, where $i \in T, j \in C$, as we already know how many 3-node motifs would form between locus $j$ and any another locus $k$ (as calculated above), we can also calculate the number of 4-node motifs involving $i, j$ by counting the 3-node motifs using the similar method. The complexity of this part would also be $O(tc^2)$. Finally, for the $W_{M_{ij}}$, where $i \in T, j \in T$, we cannot count the 3-node triangle anymore. To count the number of 4-node motifs involving $i, j$, we find out the common loci they share, and then calculate the total number of edges between these common loci. To summarize, the runtime for the whole algorithm is dominated by the construction of $W_M$, especially for the part between TF and TF. The worst case runtime would be $O(t^2c^2)$, where we have to go over all the combination of two TF and two gene loci. Note that, however, TFs only make up a small part of the nodes (about 4.5%). Also, as the network is somehow sparse, usually, we do not need to go over all the combination of nodes, which would accelerate the computation further. In addition, in the actual implementation, we use parallel computation to further speed up the process, which makes the entire algorithm quite efficient in practice.

#### A.3 Clusters are near optimal

Here we prove that the two clusters from Steps (1) and (2) described in the algorithm in the main text are near optimal when $\alpha = 4/3$ in Eq. (3). Without loss of generality, we prove that the two clusters $S$ and $\bar{S}$ of the original heterogeneous network $G$ are near optimal. By definition, $\varphi_M(S) = \varphi_M(\bar{S})$, thus we only need to show that $S$ is near optimal.

We first formally state the near optimal claim and prove it in the subsequent paragraphs. Let $\varphi_M^\star$ be the minimum of subgraph conductance over all possible sets of nodes in $G$. Then $S$ satisfies motif Cheeger inequality (Chung, 2007), i.e.,

$$\varphi_M(S) \leq 4\sqrt{\varphi_M^\star} \leq 1 \tag{7}$$

which means that $S$ is at most a quadratic factor away from the optimal cluster that achieves $\varphi_M^\star$.

We recall and define some mathematical notations. Let $N$ be the total number of nodes in $G$. Let $M$ be the subgraph with four nodes and five interactions, where the four nodes are 2 TFs and 2 genes. Four of the five interactions are interactions between TFs and genes. The fifth interaction is between the two genes. Let $V_M$ be the set of the four nodes of $M$. Let $|V_M|$ denote the cardinality of $V_M$. Here $|V_M| = 4$. Let $\mathbb{M}$ be the set of the occurrences of $M$ in $G$. Let $W_M$ denote the subgraph adjacency matrix where $[W_M]_{ij} = \sum_{M \in \mathbb{M}} \mathbb{1}(i \in V_M, j \in V_M)$. The undirected weighted network induced by $W_M$ is denoted by $G_M$. The subgraph conductance $\varphi_M(S)$ for $G$ is defined in Eq. (8) and the conductance $\varphi_{G_M}(S)$ is defined in Eq. (9):

$$\varphi_M(S) = \frac{\text{cut}_M(S, \bar{S})}{\min[\text{Vol}_M(S), \text{Vol}_M(\bar{S})]} \tag{8}$$

$$\varphi_{G_M}(S) = \frac{\text{cut}_{G_M}(S, \bar{S})}{\min[\text{Vol}_{G_M}(S), \text{Vol}_{G_M}(S)]} \tag{9}$$

First, we prove that $\text{cut}_M(S,\bar{S}) = \frac{1}{3}\text{cut}_{G_M}(S,\bar{S})$. Let $X = (x_1, x_2, \ldots, x_N)$ be the vector denoting which nodes belong to $S$. If node $i$ belongs to $S$, then $x_i = -1$. Otherwise, $x_i = 1$. Let $v_1, v_2, v_3, v_4$ be the four nodes in an occurrence of the subgraph $M \in \mathbb{M}$. We have:

$$
\begin{aligned}
\text{cut}_M(S,\bar{S}) &= \sum_{M\in\mathbb{M}} \mathbb{1}\left(|V_M \cap S| \in \{1,3\}\right) + \frac{4}{3}\sum_{M\in\mathbb{M}} \mathbb{1}\left(|V_M \cap S| = 2\right) \\
&= \sum_{M\in\mathbb{M}} \frac{6\,\mathbb{1}\left(|V_M \cap S| \in \{1,3\}\right) + 8\,\mathbb{1}\left(|V_M \cap S| = 2\right)}{6} \\
&= \sum_{M\in\mathbb{M}} \frac{6 - x_{v_1}x_{v_2} - x_{v_1}x_{v_3} - x_{v_1}x_{v_4} - x_{v_2}x_{v_3} - x_{v_2}x_{v_4} - x_{v_3}x_{v_4}}{6} \\
&= \sum_{M\in\mathbb{M}} \frac{\frac{3}{2}(x_{v_1}^2 + x_{v_2}^2 + x_{v_3}^2 + x_{v_4}^2) - (x_{v_1}x_{v_2} + x_{v_1}x_{v_3} + x_{v_1}x_{v_4} + x_{v_2}x_{v_3} + x_{v_2}x_{v_4} + x_{v_3}x_{v_4})}{6} \\
&= \frac{\frac{1}{2}x^T D_M x - \frac{1}{2}x^T W_M x}{6} \\
&= \frac{2 \times \text{cut}_{G_M}(S)}{6} \\
&= \frac{1}{3}\text{cut}_{G_M}(S).
\end{aligned}
$$

Next, we show that $\text{Vol}_M(S) = \frac{1}{3}\text{Vol}_{G_M}(S)$. Note that $|V_M| = 4$.

$$
\begin{aligned}
\text{Vol}_M(S) &= \sum_{i\in S}\sum_{M\in\mathbb{M}} \mathbb{1}(i \in V_M) \\
&= \sum_{i\in S}\sum_{M\in\mathbb{M}} \frac{1}{3}\sum_{j\in V_M} \mathbb{1}(|\{i,j\} \cap V_M| = 2) \\
&= \sum_{i\in S}\sum_{M\in\mathbb{M}} \frac{1}{3}\sum_{j=1}^{N} \mathbb{1}(|\{i,j\} \cap V_M| = 2) \\
&= \frac{1}{3}\sum_{i\in S}\sum_{j=1}^{N}\sum_{M\in\mathbb{M}} \mathbb{1}(|\{i,j\} \cap V_M| = 2) \\
&= \frac{1}{3}\sum_{i\in S}\sum_{j=1}^{N}[W_M]_{ij} \\
&= \frac{1}{3}\text{Vol}_{G_M}(S).
\end{aligned}
$$

We have $\varphi_M(S) = \varphi_{G_M}(S)$ by definitions in Eq. (8), Eq. (9), and that $\text{cut}_M(S,\bar{S}) = \text{cut}_{G_M}(S)$ and $\text{Vol}_M(S) = \text{Vol}_{G_M}(S)$.

Finally, let $\varphi^\star_{G_M}$ be the minimum of conductance over all possible sets of nodes $G_M$. Then $S$ satisfies the Cheeger inequality, i.e.,

$$\varphi_{G_M}(S) \leq 4\sqrt{\varphi^\star_{G_M}} \leq 1.$$

##### *Comparison with the proofs in Benson et al. (2016)*

In Step (2), we apply a spectral clustering method to find two sets $S$ and $\bar{S}$ in the undirected, weighted network $G_M$ that is induced by $W_M$. The spectral clustering method is the same as the method in (Benson et al., 2016) where $W_M$ is computed based on a homogeneous motif and homogeneous network. Benson et al. (2016) proved that $S$ and $\bar{S}$ are near optimal for their case. However, the results in (Benson et al., 2016) are not applicable to our situation, because our input network and motif are heterogeneous, and converting the heterogeneous network and motif to homogeneous network and motif will mis-count the occurrences of the heterogeneous motif $M$. However, our proofs follow the same strategy as the proofs in (Benson et al., 2016).

##### A.4 Properties of the identified HIMs

Here we describe the properties of HIMs that are not defined in the main text. The features are the topological structural features related to connection patterns of the genes and TFs in a given HIM within a heterogeneous network, including motif density, chromatin interaction edge density (or Hi-C edge density), and GRN edge density, which quantify the connection strength between genes/TFs in a HIM in terms of different connection patterns. All range from 0 to 1.

- 4-node motif $M$ density. It is the ratio of the number of occurrences of the motif $M$ to the total number of possible occurrences of the motif $M$ in a HIM. The maximal motif density is 1, which is achieved when every pair of genes in the cluster are connected with Hi-C interactions and every gene is regulated by each TF in the cluster. The triangle motif density used later in Supplemental Information B.2 is defined similarly.
- Hi-C edge density. It is the density of the sub-Hi-C interaction network induced by the genes in the HIM. The Hi-C edge density at 1 means that every pair of genes is connected by a chromatin interaction with O/E$>$1, where 0 means that no pair of genes is connected. Thus a higher density means that the HIM genes as a unit are more densely packed in the nucleus.
- GRN edge density. It is the density of the sub-GRN induced by the genes and TFs in the HIM. The maximal 1 is achieved when every gene is regulated by every TF in the HIM. The minimal 0 is achieved when TF-gene interaction does not exist in a HIM.

##### A.5 Collection and processing of data used in this study

In this work, we use data for five human cell types: GM12878, HeLa, HUVEC, K562, and NHEK. For Hi-C related data, including KR normalized contact frequency matrices by in-situ Hi-C and O/E contact frequency matrices were from (Rao et al., 2014). We downloaded the data from GEO with the accession number GSE63525. We calculated the A/B compartments for each chromosome using the first principal component of the O/E contact frequency matrix as the same in (Lieberman-Aiden et al., 2009). For intra-chromosomal contacts, we first filtered the genome-wide KR normalized contact matrix by only keeping intra-chromosomal contacts higher than expected values, aiming to reduce intra-chromosomal contacts due to random chromatin collisions. In addition, we used the size of compartments to control the 1D distance between genes farthest apart in HIMs. The 99-th percentile of the size of compartments is around 10Mb in 4 out of the 5 cell types. Thus we chose the 1D distance cutoff universally as 10Mb and only kept the remaining intra-chromosome contacts that connect bins within 10Mb across the cell types. Then only the top 1% inter-chromosomal contacts were kept by choosing the cutoff as the 99-th percentile of the genome-wide inter-chromosomal contacts. The remaining inter-chromosomal contacts have at least 2.17 KR normalized Hi-C contacts. Processed replication timing data with the GEO accession number GSE34399 (Hansen et al., 2010; Thurman et al., 2007) were downloaded from the UCSC Genome Browser (Rosenbloom et al., 2012).

GRN data were downloaded from (Marbach et al., 2016), where directed TF-gene interactions were inferred by simultaneously considering the TF binding motifs and gene expression level. Briefly, an interaction between a TF and a gene is called if (1) the TF has enriched binding motifs on the enhancer or promoter regions of the gene; and (2) the co-expression level between the TF and the enhancer or promoter of the gene is high.

Protein-protein interactions (PPIs) were downloaded from BioPlex2 (Huttlin et al., 2017), BioGrid (Chatr-Aryamontri et al., 2012), CORUM (Ruepp et al., 2009), and STRING (Franceschini et al., 2012). We first extracted the PPIs between the 591 TFs in the GRNs from these public sources. We then combined them into a PPI network after merging duplicated PPIs. Note that the GRNs have the same set of TF protein. Thus the PPI network is suitable for all 5 different cell types. The density of the PPI network is 0.158, which is also the expected density of a sub-PPI network of a set of randomly sampled TFs.

Essential genes in four cell lines were downloaded from (Wang et al., 2015). However, among the four cell lines, only K562 matches the cell types used in this study. The essential genes identified in K562 were only used to analyze identified HIMs in K562. Because the majority of the essential genes are shared between the 4 cancer cell lines (Wang et al., 2015), we used the union of them as the essential gene list in GM12878, HeLa, HUVEC, and NHEK cell types. The union has 2741 essential genes.

RNA-seq data with GEO accession # GSE33480 were downloaded from the ENCODE project (Consortium et al., 2012). The gene expression level quantified in FPKM value across the 5 cell types were normalized by quantile normalization then logarithm transformed by the function $\log_{10}(1+x)$.From the expression data, we constructed a list of cell type-specific genes for each cell type by the following two criteria. Given a cell type, (1) the gene expression value in the given cell type is higher than 0.1; (2) the ratio of the gene expression value in the given cell type to the median gene expression value in the other 4 cell types is higher than 2.

### B Supplementary Results

#### B.1 HIMs are robust to the parameters used to construct the heterogeneous networks

We show that the identified HIMs in the main text are robust to the parameters used to define the heterogeneous networks. In all the analysis presented in the main text, we use a cutoff at 1 for "observed over expected" (O/E) quantity to filter out intra-chromosomal Hi-C contacts when defining chromatin interaction networks. To test the robustness of HIMs, we construct another set of chromatin interaction networks by the cutoff at 2 (for O/E). The number of intra-chromosomal chromatin interactions with the cutoff at 2 is 64.6%-81.9% of the number of intra-chromosomal interactions with the cutoff at 1 across five cell types. We denote the sub-Hi-C interaction networks resulted from the cutoff at 2 as sub-Hi-C. Regarding GRNs, we also construct a sub-GRN for each cell type by only keeping the top 90% interactions with the highest scores. We then construct 3 different heterogeneous networks for each cell type as follows:

- The heterogeneous network combines the chromatin interaction network with the cutoff at 1 and the whole GRN. This is the heterogeneous network used in the main text. The heterogeneous network is referred as Hi-C + GRN.
- The heterogeneous network combines the chromatin interaction network with the cutoff at 2 and the whole GRN. The heterogeneous network is referred as sub-Hi-C + GRN.
- The heterogeneous network combines the chromatin interaction network with the cutoff at 1 and the sub-GRN. The heterogeneous network is referred to as Hi-C + sub-GRN.

We apply MOCHI with the 4-node motif $M$ to each of the 3 heterogeneous networks of each cell type. We use adjusted Rand index to quantify the similarities on gene memberships between the HIMs from two different heterogeneous networks. For example, if the assignment of genes to HIMs are identical between two sets of HIMs, then adjusted Rand index would be 1. We use hierarchical clustering to group the sets of HIMs with similar adjusted Rand index. We found that the sets of HIMs from the heterogeneous networks of the same cell type are much more similar to each other than to the sets of HIMs from the other cell types. Hierarchical clustering produces five major clusters (Fig. S10). Each cluster contains 3 different heterogeneous networks in the same cell type. Overall, the result suggests that the HIMs are less sensitive to the parameters used in constructing the chromatin interactome and GRNs.

#### B.2 Justification of the 4-node motif *M*

To justify the choice of the motif $M$, we compared it with two different types of motifs. One is a triangle motif with 3 nodes (Fig. S11A), where two of them are genes with a chromatin interaction and the third node is a TF that regulates both the genes. The triangle motif does not explicitly encode co-regulation between TF proteins. Another motif is a bifan motif with 4 nodes (Fig. S11A), where two nodes are TF proteins that co-regulate two genes but there is no chromatin interaction between the two genes. The bifan motif does not explicitly encode spatial proximal relationship between genes. For bifan motif, we applied MOCHI with the bifan on the GRN and then split the identified HIMs by chromosome number.

We found that the motif $M$ and triangle motif are better than the bifan motif in terms of identifying the clusters (Fig. S11B). Compared to the HIMs identified by the bifan motif, the HIMs by the motif $M$ and triangle motif have higher Hi-C edge density, higher triangle density, higher motif $M$ density. Moreover, the genes in a HIM are closer to each other in the 1D sequence space, although the HIMs by the bifan motif have a smaller number of genes as compared to the HIMs by the motifs $M$ and triangle. This result highlights that the chromatin interaction between the two target genes in a motif is important to capture spatial proximity between the genes in HIMs.

In addition, we found that our motif $M$ is better than the triangle motif. We comprehensively compared the identified HIMs by the motif $M$ and the triangle motif. The HIMs identified by the two motifs have similar numbers of genes as the median numbers of genes are equal in the 4 out of 5 cell types. A similar pattern is observed on the Hi-C edge density. Specifically, the density is only significantly different in two cell types: GM12878 and NHEK ($p \leq 0.04$), although the difference is small (the median density is 0.015 in GM12878 and 0.028 in NHEK) (Fig. S11B). However, the identified HIMs by the two motifs are very different in other features. Compared to the HIMs identified by the triangle motif, the HIMs identified by the 4-node motif $M$ have much higher numbers of TFs ($p \leq$ 2.45e-21). The difference in the median number of TFs ranges from 4 to 8 (Fig. S12). Even though the HIMs by the 4-node motif $M$ have higher number TFs, they have comparable numbers of genes as compared to the HIMs from the triangle. They also have much higher triangle density and much higher 4-node motif $M$ density ($p \leq$2.68e-02 and $p \leq$1.43e-03, respectively; Fig. S11B). The HIMs from $M$ also have a higher proportion of genes in the A compartment in 4 cell types ($p \leq$2.18e-02), are much earlier replicated ($p \leq$6.74e-03), and have smaller replication timing coefficient of variation ($p \leq$4.38e-05) (Fig. S12).

Next, we compared the features after adjusting the number of TFs and the number of genes of the identified HIMs. The HIMs by the 4-node motif have much higher numbers of TFs than the HIMs by the triangle motif. The number of genes is slightly different in some cell types. Since the features could be biased to the number of TFs and the number of genes, we compared the features by adjusting the number of genes and the number of TFs in HIMs by a linear regression model:

$$Y = \beta_0 + \beta_1 \times \text{\# TFs} + \beta_2 \times \text{\# genes} + \beta_3 \times \mathbb{1}_{\text{motif}}, \quad (10)$$

where $Y$ is a given feature, $\mathbb{1}_{\text{motif}} = 1$ if the HIM is identified with the 4-node motif $M$, $\mathbb{1}_{\text{motif}} = 0$ if the HIM is identified with the triangle motif, and $\hat{\beta}_3$ indicates the averaged difference in the feature $Y$ between the HIMs identified by the two motifs after adjusting the number of TFs and the number of genes in the HIMs. Specifically, a positive $\hat{\beta}_3$ means that HIMs with the 4-node motif is higher in the feature $Y$ than the HIMs with the triangle motif. On the other hand, a negative $\hat{\beta}_3$ means lower $Y$ in HIMs with the 4-node motif $M$. The detailed $\hat{\beta}_3$ for the features are reported in the Table S4. Overall, the differences are still significant after adjusting the number of TFs and the number of genes. Take together, the 4-node motif $M$ is better than the triangle motif in identifying HIMs.

#### B.3 HIMs share similar connections with 3D genome features across cell types

We found that the HIMs in 5 cell types in this study share similar connections with 3D genome organization features, such as A/B compartments, TADs, and loops. We looked at the genomic regions of each HIM that is the smallest genomic block containing the transcription start sites of the genes in the HIM. The median size of the genomic regions of the HIMs ranges from 4.9Mb in NHEK to 8Mb in GM12878 (Table S2), comparable to the size of A/B compartments (median size is 5Mb). The median numbers of TADs in the genomic regions of the HIMs are 3-4 in different cell types (Table S2). The genomic regions of the HIMs have, on average, 7 chromatin loops in GM12878 and 2-4 loops in the other cell types (Table S2). The HIMs in GM12878 involve a higher number of loops, which perhaps is due to the fact that GM12878 has at least 60% more detected loops than the other cell types possibly due to higher sequencing depth (Rao et al., 2014).

**Table S1:** Summary of the input heterogeneous networks and the identified HIMs across five cell types. 'Overlapping HIMs (%)' is the proportion of identified HIMs that share TFs with other HIMs. 'Genes in HIMs (%)' represents the proportion of genes in a heterogeneous network that are assigned to HIMs.

| | | GM12878 | HeLa | HUVEC | K562 | NHEK |
|---|---|---|---|---|---|---|
| Input | TFs | 591 | 591 | 591 | 591 | 591 |
| | Genes | 11,627 | 12,036 | 11,927 | 12,391 | 12,161 |
| | TF→gene | 1,078,893 | 998,174 | 828,303 | 1,119,395 | 814,017 |
| | Gene−gene | 337,036 | 164,007 | 184,866 | 253,218 | 139,385 |
| Output | HIMs | 650 | 806 | 773 | 802 | 664 |
| | Overlapping HIMs (%) | 72.8 | 74.7 | 71.9 | 74.7 | 79.4 |
| | Genes in HIMs (%) | 69.1 | 77.2 | 75.3 | 76.5 | 62.1 |

**Table S2:** Statistics of the identified HIMs across five cell types.

| | GM12878 | HeLa | HUVEC | K562 | NHEK |
|---|---|---|---|---|---|
| Median TF number | 9 | 17 | 17 | 15 | 14 |
| Median gene number | 9 | 9 | 9 | 9 | 9 |
| Median loop number | 7 | 2 | 2 | 4 | 2 |
| Median TAD number | 4 | 3 | 3 | 4 | 3 |
| Median 1D distance between The farthest apart genes (Mb) | 8 | 5.4 | 6.1 | 7.2 | 4.9 |
| # of HIMs inherited TFs | 451 | 599 | 546 | 596 | 497 |
| Median proportion of inherited TFs | 28.6 | 27.8 | 24.6 | 25.0 | 28.0 |

**Table S3:** The dynamics of chromatin interaction networks and GRNs across 5 cell types. Cells in the table are numbers/proportions of interactions that exist in the corresponding number of cell type in each column. For example, column '1' corresponds to the interactions that only exist in one cell type. Column '5' corresponds to the interactions that exist in all 5 different cell types. Overall, a large proportion of the edges in the GRNs and chromatin interaction networks only exist in one cell type.

| | Type | 1 | 2 | 3 | 4 | 5 |
|---|---|---|---|---|---|---|
| Hi-C networks | # interactions | 755,574 | 242,833 | 86,790 | 36,945 | 23,442 |
| | % of interactions | 66.00 | 21.20 | 7.60 | 3.20 | 2.00 |
| GRNs | # interactions | 637,950 | 457,289 | 309,733 | 234,033 | 389,722 |
| | % of interactions | 31.40 | 22.50 | 15.30 | 11.50 | 19.20 |

**Table S4:** Comparison between the identified HIMs by the 4-node motif $M$ and the triangle motif while adjusting the numbers of TFs and genes in the HIMs by the linear regression model $Y = \beta_0 + \beta_1 \times$ # TFs $+ \beta_2 \times$ # genes $+ \beta_3 \times \mathbb{1}_{\text{motif}}$, where $Y$ is a continuous feature, $\mathbb{1}_{\text{motif}} = 1$ if a HIM is identified by the 4-node motif $M$ and 0 otherwise, $\hat{\beta_3} \geq 0$ means that the HIMs identified by the 4-node motif $M$ have higher $Y$ than the HIMs identified by the triangle motif after adjusting the numbers of TFs and genes. P-value is computed for the hypothesis that $\beta_3 \neq 0$. The features with P-value $< 0.05$ across 5 cell types are highlighted with bold font.

| $Y$ | GM12878 | | HeLa | | HUVEC | | K562 | | NHEK | |
|---|---|---|---|---|---|---|---|---|---|---|
| | $\hat{\beta_3}$ | P value | $\hat{\beta_3}$ | P value | $\hat{\beta_3}$ | P value | $\hat{\beta_3}$ | P value | $\hat{\beta_3}$ | P value |
| Hi-C edge density | 0.005 | 5.86e-01 | 0.012 | 1.55e-01 | 0.02 | 1.51e-02 | -0.001 | 9.03e-01 | 0.026 | 3.02e-03 |
| **Triangle density** | 0.11 | 9.9e-17 | 0.071 | 3.19e-10 | 0.074 | 6.04e-11 | 0.056 | 4.45e-07 | 0.077 | 1.43e-11 |
| **4-node motif $M$ density** | 0.15 | 1.93e-22 | 0.082 | 1.83e-11 | 0.089 | 6.25e-13 | 0.074 | 1.35e-09 | 0.09 | 6.91e-13 |
| % of genes in A compartment | 0.038 | 3.61e-04 | 0.036 | 8.4e-03 | 0.022 | 1.47e-01 | 0.033 | 7.81e-04 | 0.015 | 2.89e-01 |
| **Mean replication timing** | 2.76 | 8.87e-07 | 2.269 | 1.36e-08 | 2.485 | 2.59e-06 | 2.527 | 2.13e-07 | 2.673 | 1e-10 |
| **Replication timing CV** | -0.04 | 3.27e-13 | -0.029 | 6.88e-09 | -0.032 | 8.82e-10 | -0.029 | 2.03e-09 | -0.029 | 9.36e-12 |

**Table S5:** The top GO terms or pathways that are enriched in the genes that are assigned to HIMs consistently or in a cell type-specific manner. The number of genes in each category is shown in Fig. 4A.

| | **GO term/Pathway** | **Count** | **Fold Enrichment** | ***P* value** |
|---|---|---|---|---|
| **Constitutive genes** | chromosome organization | 371 | 1.50 | 5.7e-19 |
| | macromolecular complex subunit organization | 653 | 1.30 | 1.2e-12 |
| | regulation of gene expression, epigenetic | 105 | 1.90 | 3.4e-12 |
| | RNA processing | 287 | 1.40 | 4.1e-12 |
| | nucleosome organization | 74 | 2.10 | 4.2e-12 |
| | DNA conformation change | 104 | 1.80 | 2.2e-11 |
| | mRNA processing | 163 | 1.60 | 9.3e-11 |
| | protein-DNA complex subunit organization | 98 | 1.80 | 2.0e-10 |
| | DNA packaging | 77 | 1.90 | 5.2e-10 |
| | mRNA metabolic process | 213 | 1.40 | 2.5e-9 |
| | RNA splicing | 139 | 1.60 | 3.4e-9 |
| | protein localization to organelle | 268 | 1.40 | 4.2e-9 |
| | intracellular transport | 436 | 1.30 | 6.5e-9 |
| **GM12878 specific genes** | regulation of lymphocyte activation | 28 | 3.40 | 6.5e-8 |
| | regulation of T cell activation | 24 | 3.80 | 7.4e-8 |
| | regulation of leukocyte cell-cell adhesion | 24 | 3.60 | 1.8e-7 |
| | T cell activation | 28 | 3.00 | 5.5e-7 |
| **HeLa specific genes** | cell development | 74 | 1.50 | 4.8e-4 |
| | cell-cell signaling | 57 | 1.60 | 5.6e-4 |
| **K562 specific genes** | phospholipase C-activating G-protein coupled receptor signaling pathway | 11 | 4.80 | 8.5e-5 |
| | reproduction | 55 | 1.70 | 1.3e-4 |
| | G-protein coupled receptor signaling pathway | 33 | 2.00 | 1.7e-4 |
| **NHEK specific genes** | keratinocyte differentiation | 23 | 10.30 | 2.5e-16 |
| | skin development | 30 | 6.60 | 1.2e-15 |
| | keratinization | 16 | 18.30 | 2.0e-15 |
| | peptide cross-linking | 16 | 17.00 | 7.0e-15 |
| | epidermis development | 32 | 5.60 | 2.6e-14 |

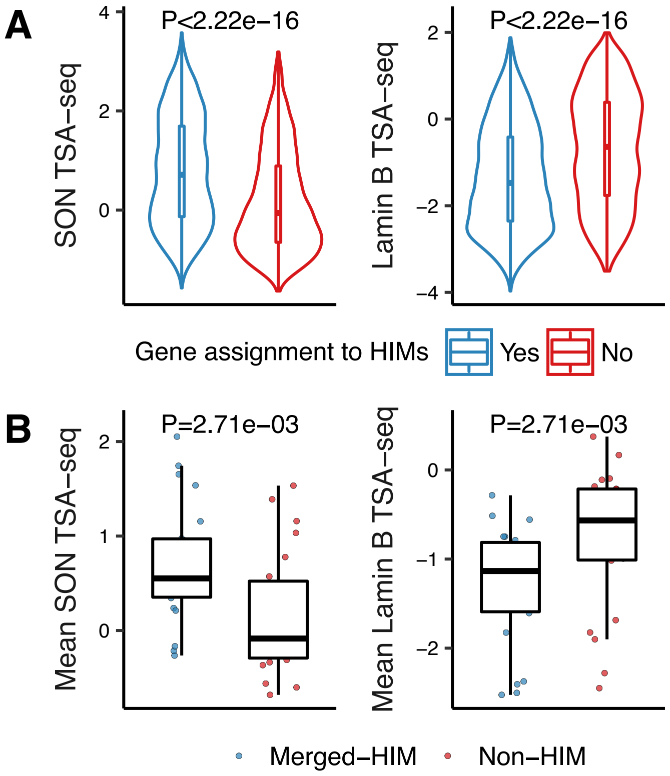


**Figure S1:** Genes assigned to HIMs are closer to nuclear speckles as compared to the genes in the heterogeneous network that are not assigned to HIMs in K562. **(A)** Violin plots show the distributions of TSA-seq scores of the two sets of genes. **(B)** Boxplots show the distributions of mean TSA-seq scores of the merged-HIM clusters and non-HIM clusters. Here we merged the genes assigned to HIMs on the same chromosome into one cluster and called it a merged-HIM cluster. Similarly, we merged the genes not assigned to HIMs on the same chromosome into one cluster and called it a non-HIM cluster.

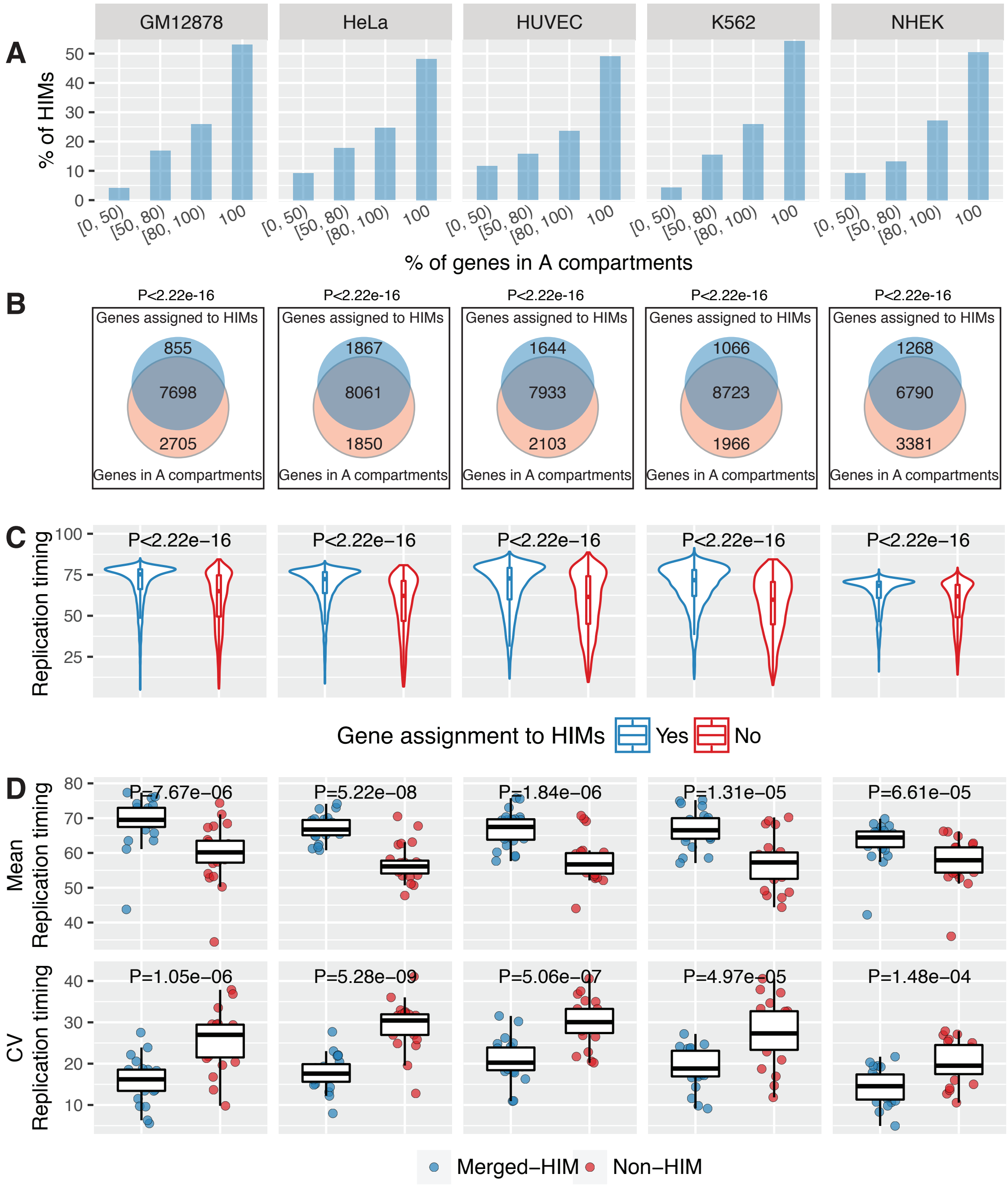


**Figure S2:** HIMs consistently have spatial location preferences as compared to non-HIMs in five cell types. Rows correspond to the spatial location features. Columns correspond to the cell types. **(A)** Barplot shows the distribution of HIMs with a varied proportion of genes that are in A compartment. **(B)** Venn diagram shows that the genes assigned to HIMs, as a whole, are enriched with the genes in A compartment. **(C)** Boxplots compare the replication timing of the genes that are assigned to HIMs against the genes that are not assigned to HIMs. **(D)** Boxplots show the mean and coefficient of variation (CV) of replication timing of the genes in merged-HIMs or non-HIMs. Each dot represents a merged-HIM or non-HIM. A lower CV means that the genes in a cluster have a lower variability in replication timing thus they are more likely to be replicated synchronously.

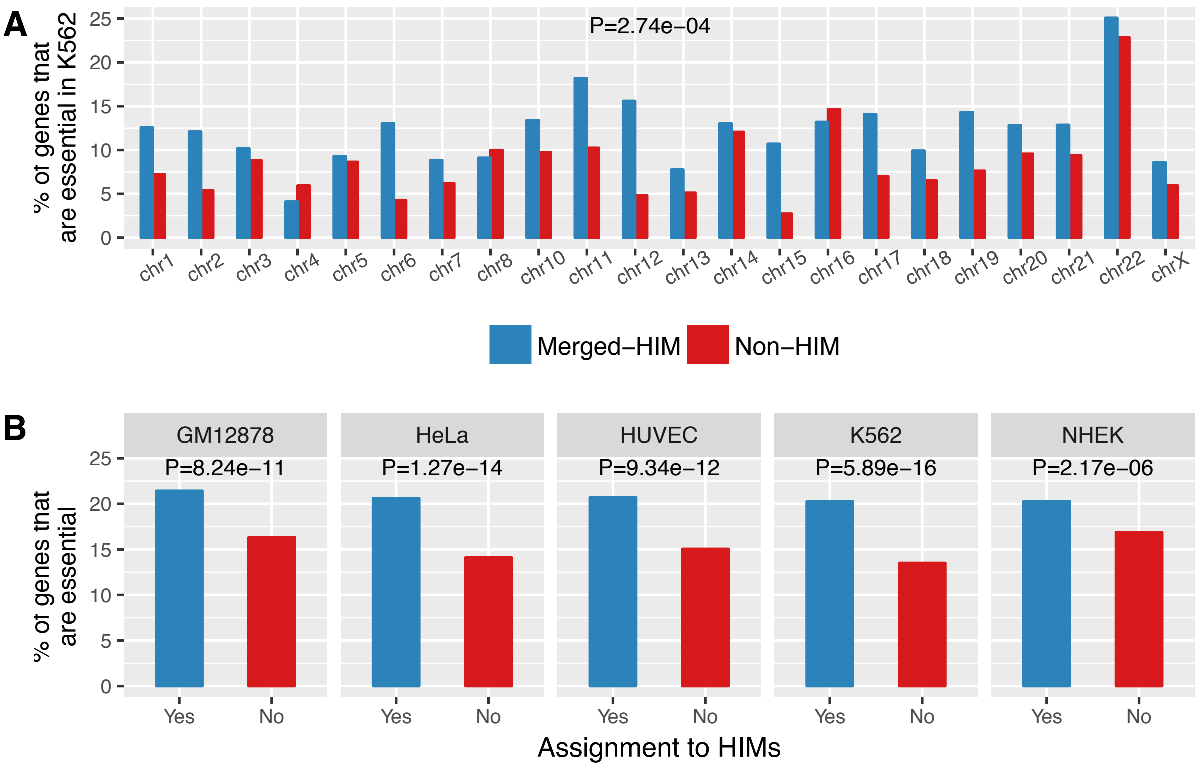


**Figure S3:** HIMs have more essential genes across cell types. **(A)** Barplots show the proportions of genes that are K562 essential genes in the genes assigned to HIMs and the genes not assigned to HIMs. **(B)** Barplots show the proportions of K562 essential genes in merged-HIMs and non-HIMs across the chromosomes. The P-value is computed by the paired two-sample Wilcoxon rank-sum test. **(C)** Barplots show the proportions of essential genes in the genes assigned to HIMs and the genes not assigned to HIMs. P-value is computed by the Chi-squared test of independence. The proportions of the essential genes in merged-HIMs and non-HIMs are similar to **(A)** thus are not shown here.

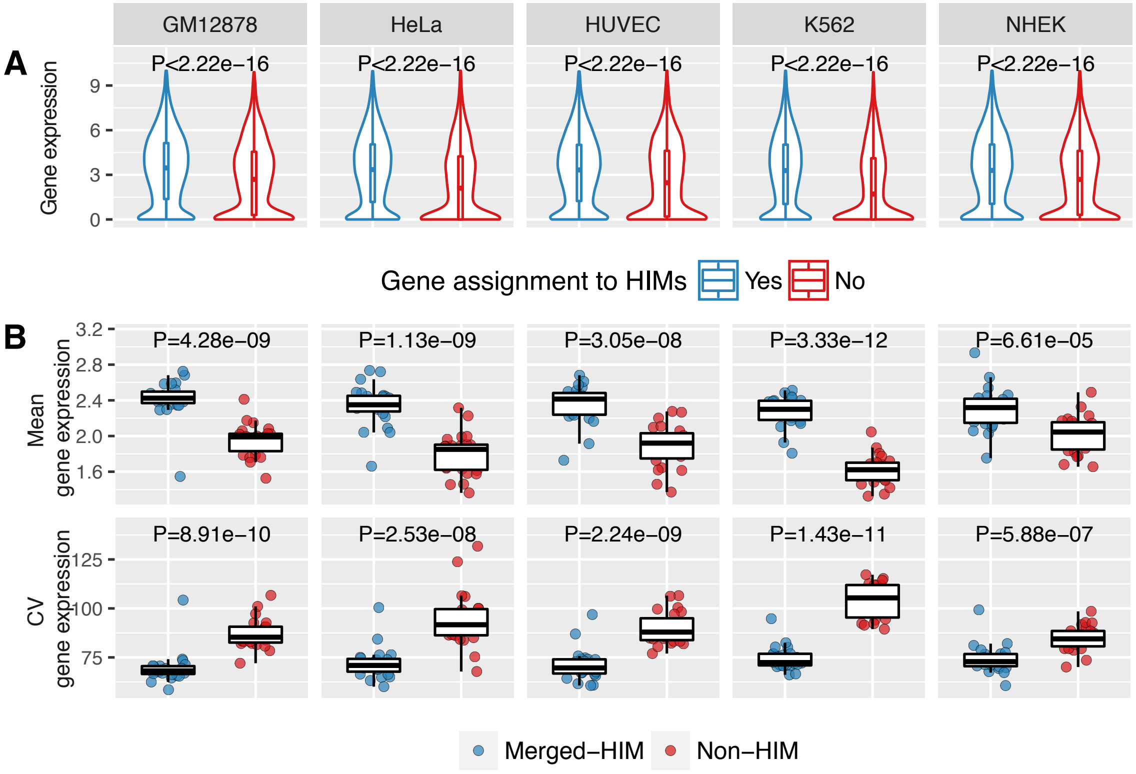


**Figure S4:** The genes assigned to HIMs express at higher levels than the genes not assigned to HIMs across cell types. **(A)** Violin plots show that the genes assigned to HIMs, as a whole, are more expressed than the genes not assigned to HIMs. **(B)** Boxplots show that the merged-HIMs have higher mean and lower CV of expression level than the non-HIMs.

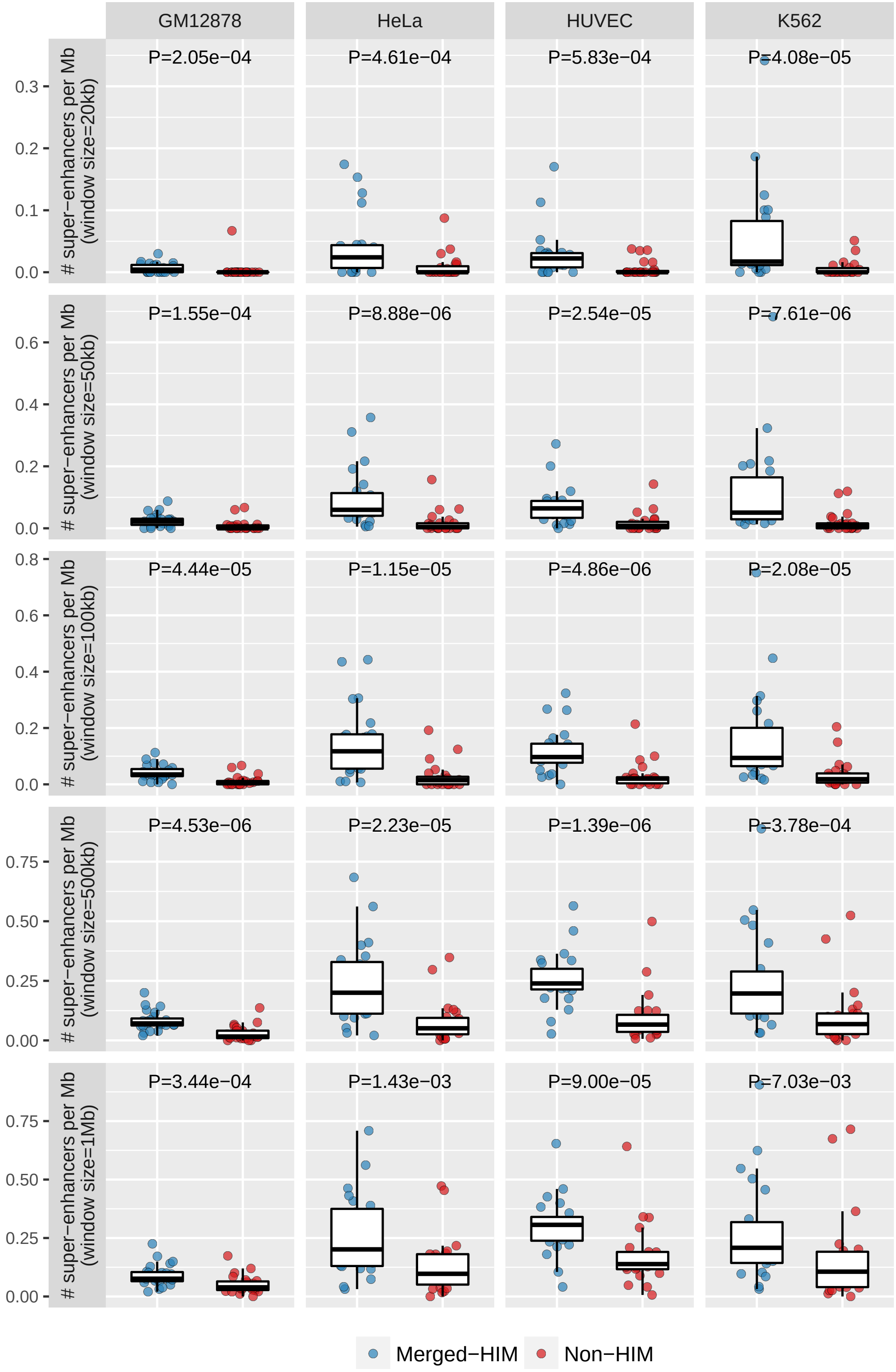


**Figure S5:** HIMs are enriched with super-enhancers and this observation is robust to the window size. The window size is used to define the genes that are close to a given super-enhancer. The window size ranges from 20kb to 1Mb. The distribution of super-enhancers in NHEK is missing due to lack of super-enhancer data in NHEK.

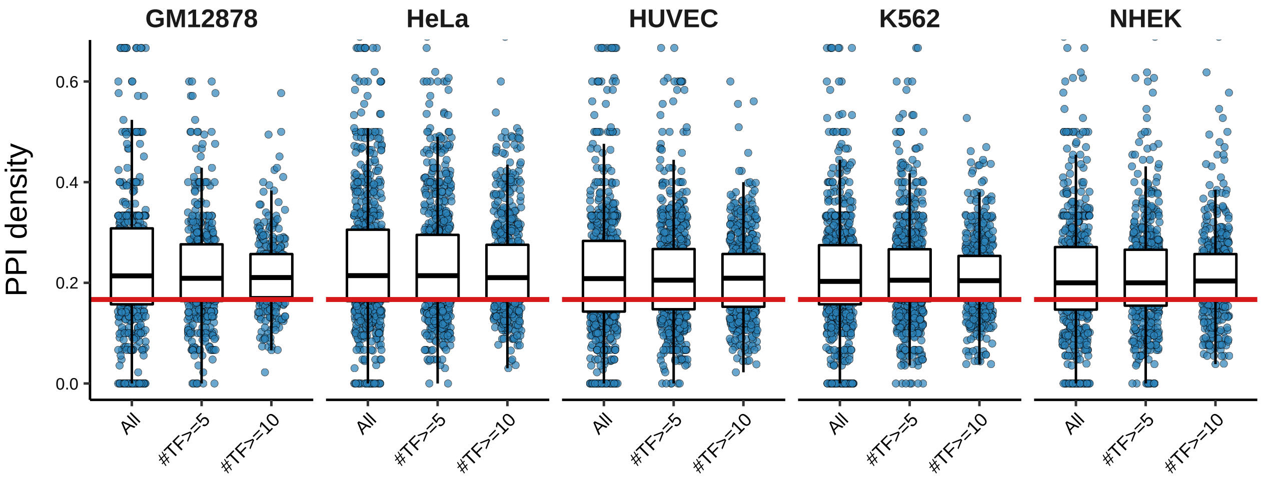


**Figure S6:** TFs in HIMs are enriched with protein-protein interactions (PPIs). Boxplots show the distribution of the sub-PPI network density across HIMs and subsets of HIMs with at least $n$ TFs, $n = 5, 10$. Here for each HIM, we computed the density of the sub-PPI network induced by the TFs in the HIM from the PPI network based on 591 TF proteins used in this study. The medians are all higher than the expected density (0.158, red line) of the sub-PPI networks induced by randomly sampled TFs ($p$<2.22e-16).

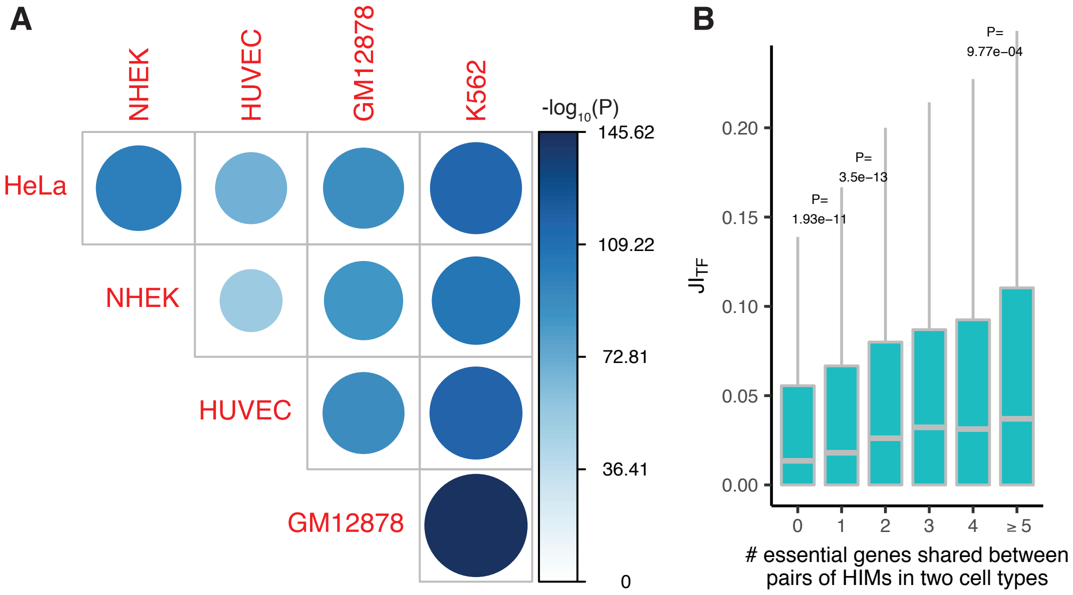


**Figure S7:** HIM comparisons regarding genes and TFs across cell types. **(A)** Heatmap shows the level of significance in overlapping between the genes assigned to HIMs in two different cell types. GM12878 and K562 have the highest overlap. Statistical significance is evaluated by the hypergeometric test. **(B)** Boxplots show the distribution of Jaccard index on the TFs of paired HIMs with different numbers of shared essential genes.

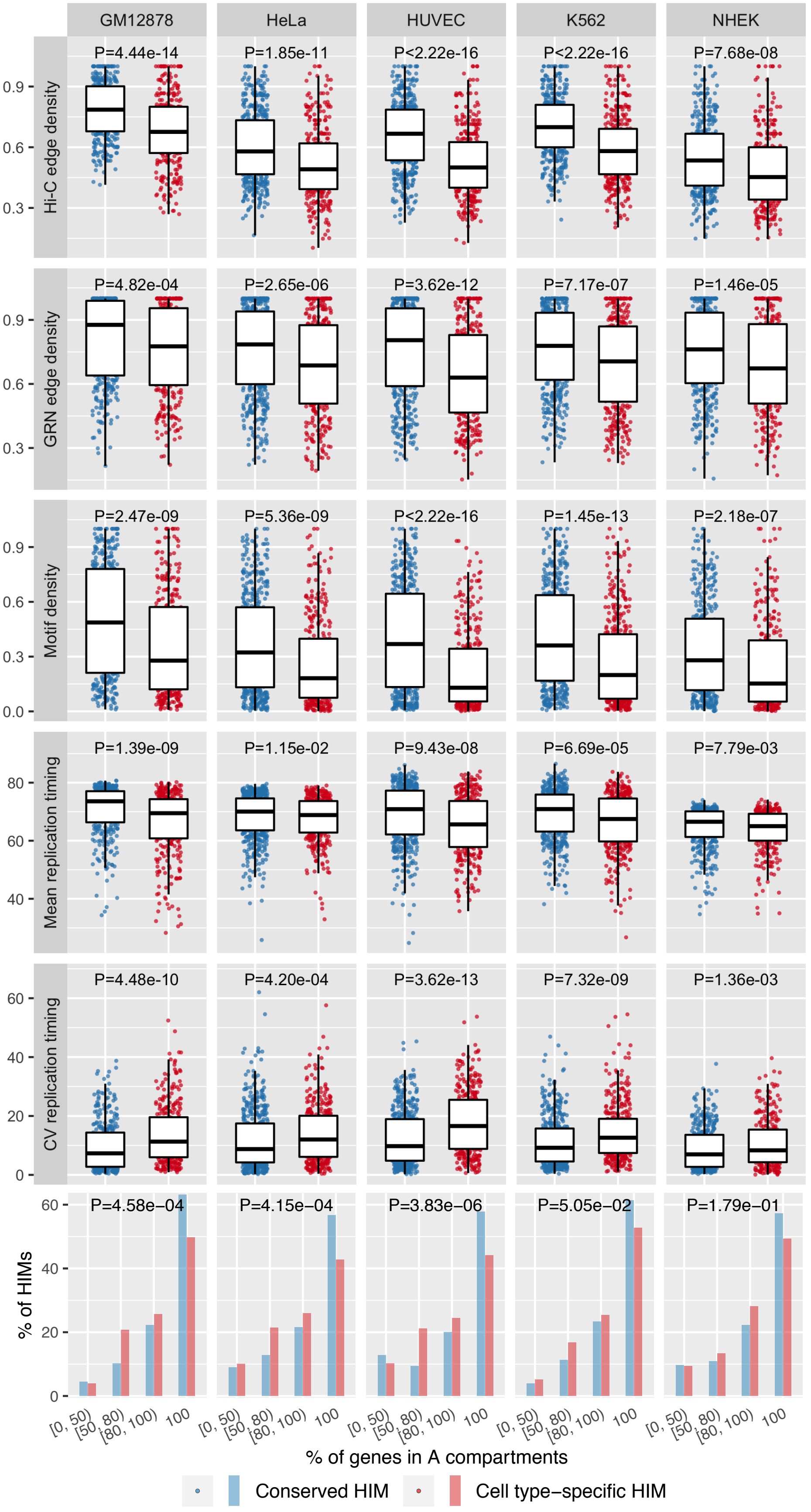


**Figure S8:** Conserved and cell type-specific HIMs have distinct spatial location features.

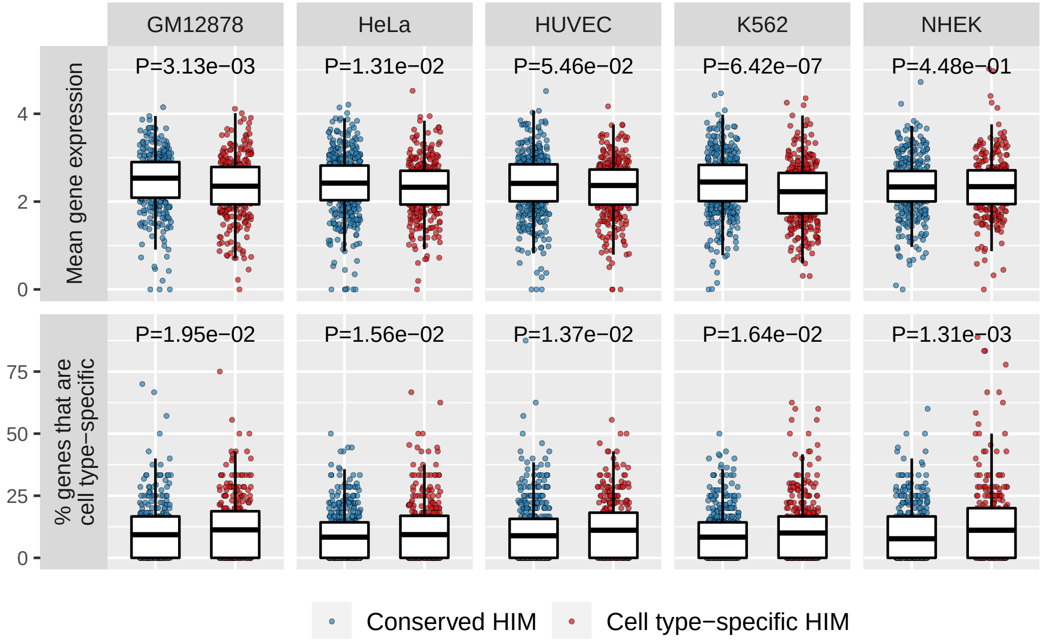


**Figure S9:** Functional differences and similarities between conserved and cell type-specific HIMs. Conserved HIMs have higher average gene expression in 3 cell types (first row). On the other hand, cell type-specific HIMs tend to have a higher proportion of cell type-specific genes in all 5 cell types (second row).

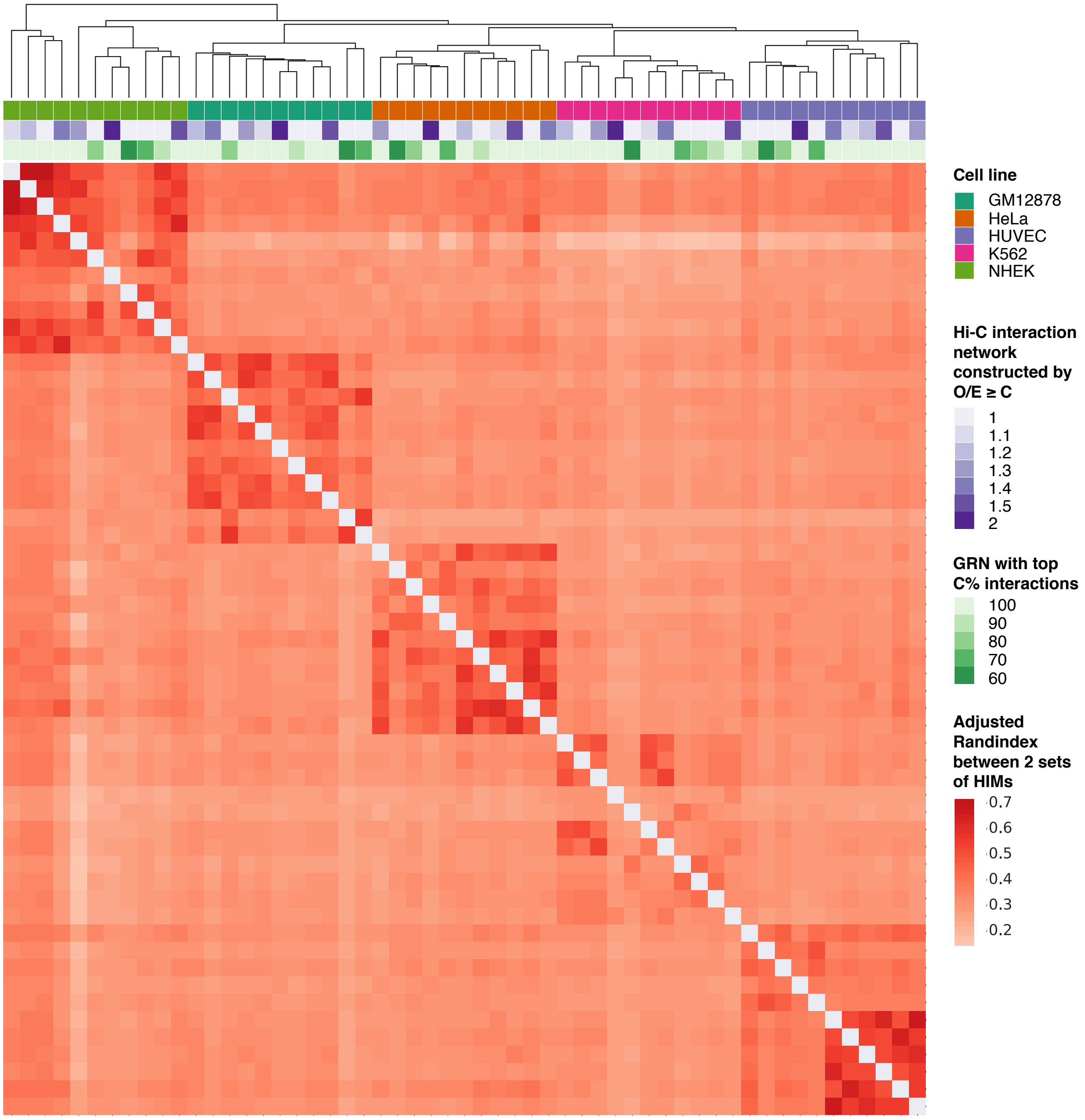


**Figure S10:** HIMs are robust to the input heterogeneous networks. For each cell type, there are 3 heterogeneous networks: one used in the main text and two sub-heterogeneous networks derived from different combinations of Hi-C interaction networks and GRNs. A sub-Hi-C represents a sub-Hi-C interaction network with interactions satisfying O/E$\geq 2$. The sub-GRN represents a sub-GRN with interactions having top 90% scores. The heterogeneous network (Hi-C + GRN) is used in the main text. The other two (sub-Hi-C + GRN, Hi-C + sub-GRN) are sub-heterogeneous networks. Adjusted Rand index is used to quantify similarities on gene memberships between two different sets of HIMs that are resulted from two different heterogeneous networks. Then hierarchical clustering on adjusted Rand index is used to cluster the sets of HIMs. Overall, the result shows that the sets of HIMs from the same cell type are in the same cluster thus are much more similar as compared to the sets of HIMs from different cell types.

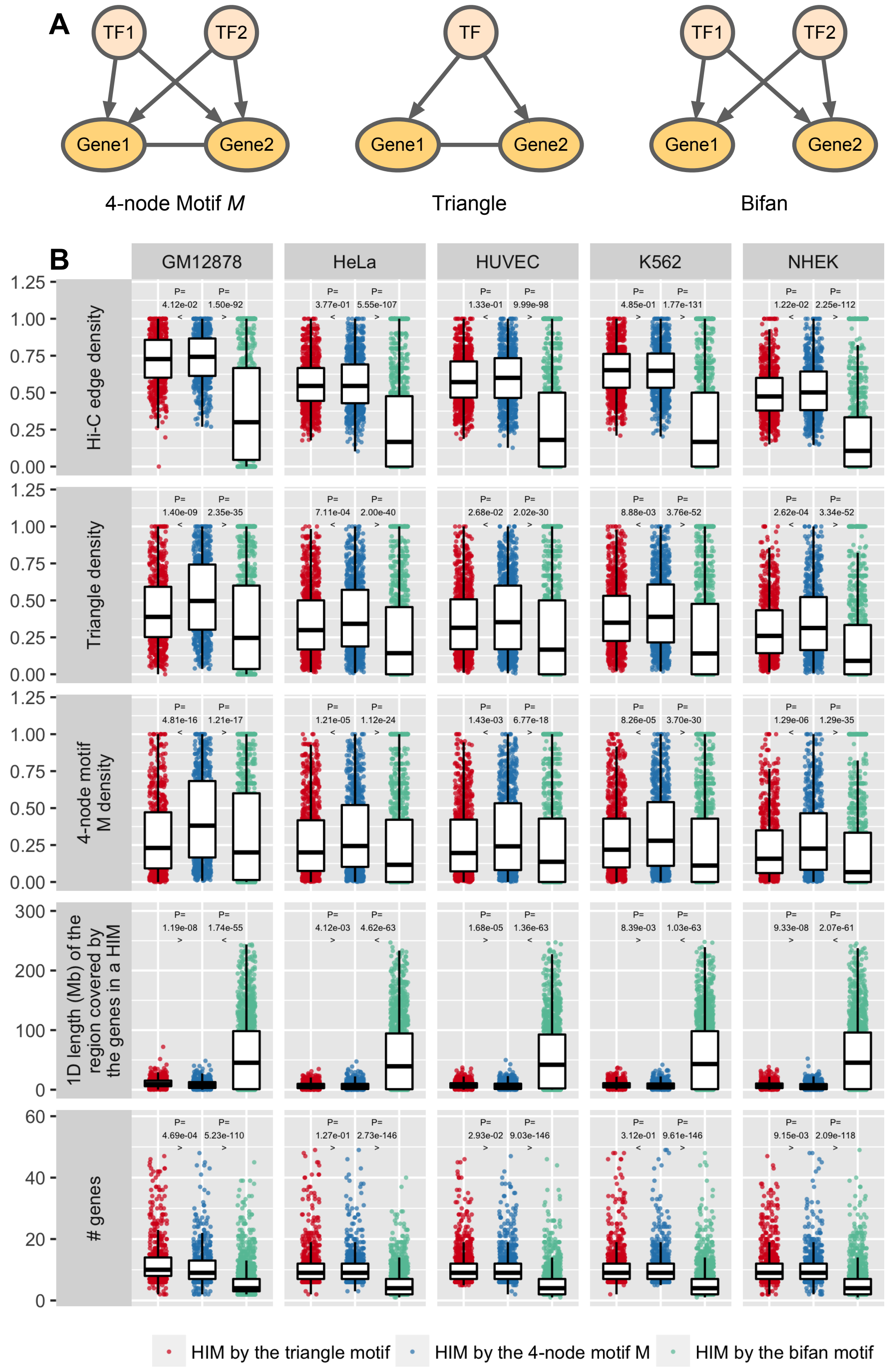


**Figure S11:** Comparison of the identified HIMs between the 4-node motif $M$, a triangle motif, bifan motif across 5 cell types. **(A)** Illustration of the 3 different motifs. **(B)** Comparison between the HIMs identified by the 3 different motifs. Rows correspond to the features. Columns correspond to the cell types. Each dot represents a HIM.

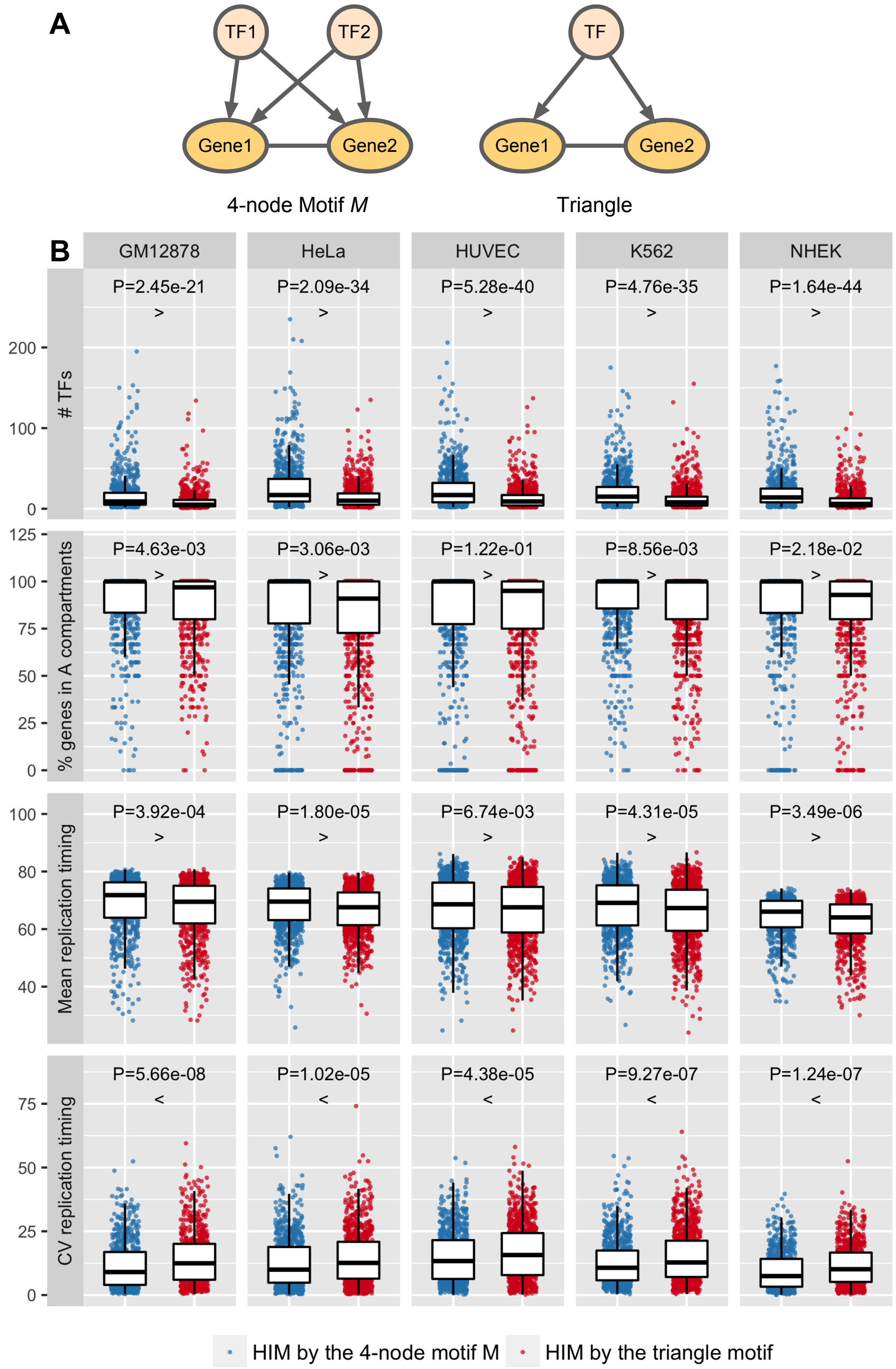


**Figure S12:** Comparison of identified HIMs between the 4-node motif $M$ and the triangle motif across the 5 cell types. Where the triangle motif has 3 nodes **(A)**. Two of them are genes connected by a chromatin interaction and co-regulated by a TF. **(B)** Rows correspond to the features. Columns correspond to the cell types. Each dot represents a HIM identified either by the 4-node motif $M$ or the triangle motif. The patterns are consistently observed after adjusting the numbers of TFs and genes in HIMs by a linear regression model (Table S4).
